# Distinct Single-cell Immune Ecosystems Distinguish True and *De Novo* HBV-related Hepatocellular Carcinoma Recurrences

**DOI:** 10.1101/2022.06.02.494526

**Authors:** Shuling Chen, Cheng Huang, Guanrui Liao, Huichuan Sun, Yubin Xie, Jianping Wang, Minghui He, Huanjing Hu, Zihao Dai, Xiaoxue Ren, Xuezhen Zeng, Qianwen Zeng, Guopei Zhang, Changyi Liao, Wenxuan Xie, Shunli Shen, Shaoqiang Li, Sui Peng, Dongming Kuang, Qiang Zhao, Dan G. Duda, Ming Kuang

**Affiliations:** Institute of Precision Medicine, The First Affiliated Hospital, Sun Yat-sen University, Guangzhou, 510080, China; Institute of Diagnostic and Interventional Ultrasound, The First Affiliated Hospital, Sun Yat-sen University, Guangzhou, 510080, China; Department of Liver Surgery and Transplantation, Liver Cancer Institute and Zhongshan Hospital, Fudan University, Shanghai, 200032, China; Department of Liver Surgery, Center of Hepato-Pancreato-Biliary Surgery, The First Affiliated Hospital, Sun Yat-sen University, Guangzhou, 510080, China; Department of Gastroenterology and Hepatology, The First Affiliated Hospital, Sun Yat-sen University, Guangzhou, 510080, China; Department of Oncology, Cancer Center, The First Affiliated Hospital, Sun Yat-sen University, Guangzhou, 510080, China; Clinical Trials Unit, The First Affiliated Hospital, Sun Yat-sen University, Guangzhou, 510080, China; State Key Laboratory of Oncology in South China, Cancer Center, MOE Key Laboratory of Gene Function and Regulation, School of Life Sciences, Sun Yat-sen University, Guangzhou, 510080, China; Organ Transplant Center, The First Affiliated Hospital, Sun Yat-sen University, Guangzhou, China; Guangdong Provincial Key Laboratory of Organ Donation and Transplant Immunology, Guangzhou, China; Steele Laboratories for Tumor Biology, Department of Radiation Oncology, Massachusetts General Hospital and Harvard Medical School, Boston, Massachusetts, MA 02478, USA

**Keywords:** HCC recurrence, tumor immune microenvironment, scRNA-seq

## Abstract

Revealing differential tumor immune microenvironment (TIME) characteristics between true versus *de novo* hepatocellular carcinoma (HCC) recurrence could help optimal development and use of immunotherapies. Here, we studied the TIME of recurrent HBV-related HCCs by 5’and VDJ single-cell and bulk RNA-sequencing, flow cytometry, and multiplexed immunofluorescence. Analyses of mutational profiles, evolutionary trajectories, and clonal architecture using whole-exome sequencing identified *de novo* versus true recurrences, some of which occurred before clinical diagnosis. The TIME of truly recurrent HCCs was characterized by an increased abundance in KLRB1^+^CD8^+^ T cells with memory phenotype and low cytotoxicity. In contrast, we found an enrichment in cytotoxic and exhausted CD8^+^ T cells in the TIME of *de novo* recurrent HCCs. Transcriptomic and interaction analyses showed an upregulated GDF15 expression level on HCC cells in proximity to dendritic cells, which may have dampened antigen presentation and inhibited anti-tumor immunity in the TIME of truly recurrent lesions. In contrast, we found that myeloid cells’ crosstalk with T cells mediated T cell exhaustion and immunosuppression in the TIME of *de novo* recurrent HCC. In conclusion, our results support genomic diagnosis and immune profiling for guiding immunotherapy implementation based on the type of HCC recurrence and TIME.

**Highlights:**

1. Truly recurrent lesions are seeded before primary tumor diagnosis, and that *de novo* cancer can occur earlier than the clinically used 2-year limit.
2. ScRNA-seq unravels distinct immune ecosystems in true versus *de novo* HCC recurrences, highlighting the need for different immunotherapy strategies for two types of HCC recurrence.
3. CD8^+^ T cells in *de novo* recurrence displayed cytotoxic and exhausted phenotypes while those in truly recurrent lesions showed a memory phenotype with weak cytotoxicity.
4. HCC cells expressing the inhibitory molecule GDF15 were in the proximity of DCs only in truly recurrent lesions.
5. High GDF15 expression level was associated with truly recurrent HCC and worse prognosis.

**Graphical abstract:** 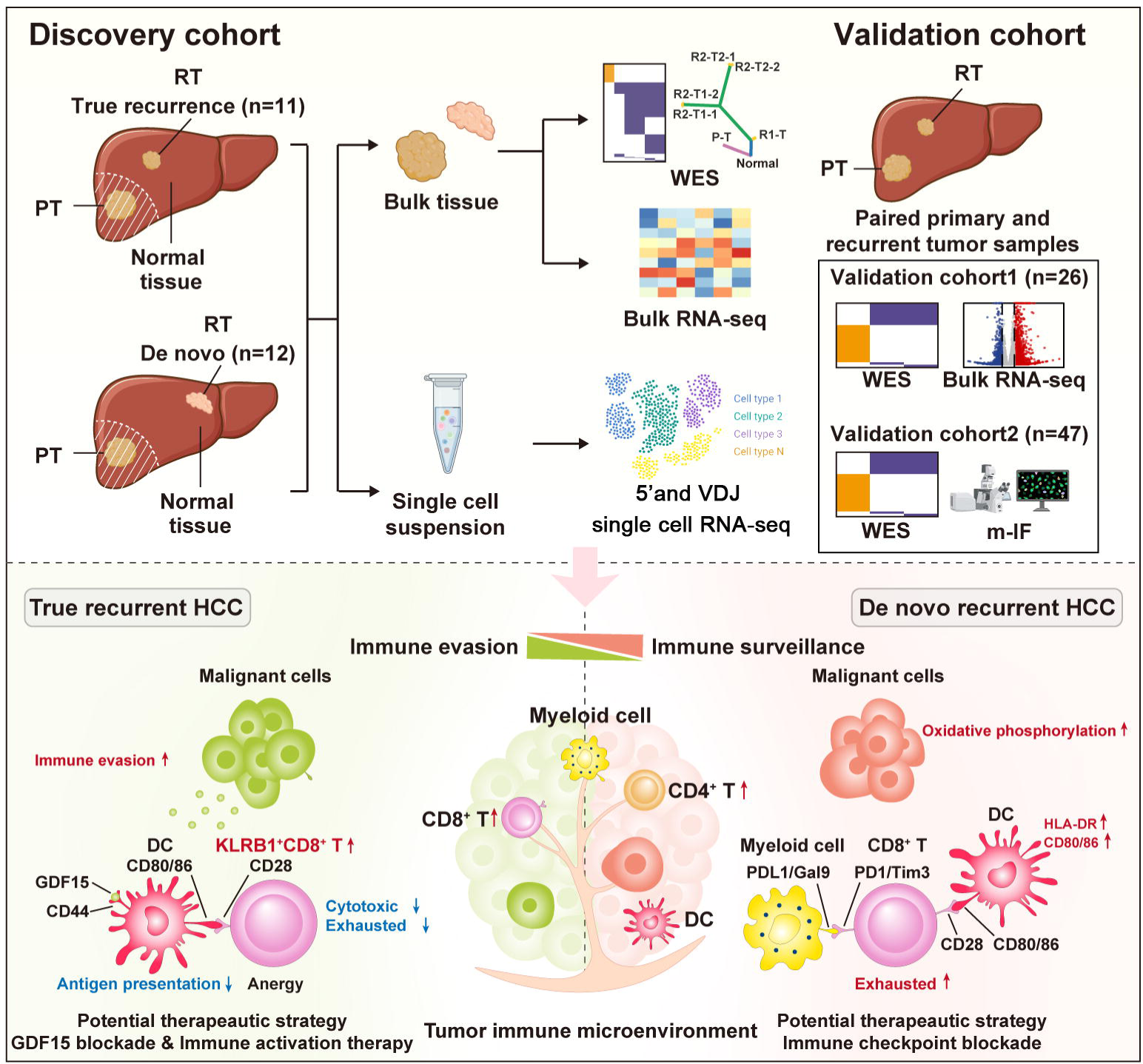

## Introduction

Hepatocellular carcinoma (HCC) is one of the most lethal cancers in the world with a 5-year survival of only 18% (Villanueva et al., 2019). The high recurrence rate after resection (70% at 5 years) is a major obstacle to long-term survival in HCC (Hasegawa et al., 2013). Integration of immunotherapies has led a paradigm shift in advanced HCC treatment, but only a minority of patients experience clinical benefits (Llovet et al., 2008; El-Khoueiry et al., 2017; Finn et al., 2020). A better understanding of the tumor immune microenvironment (TIME) of recurrent HCCs is critical to develop strategies for prevention and personalized use of immunotherapies at early stages in this aggressive disease.

HCC patients may experience true tumor recurrences (caused by primary tumor dissemination) or *de novo* cancers (newly arising from the damaged liver parenchyma as most patients also suffer from liver damage or cirrhosis) (European Association for the Study of the Liver, 2018; Chen et al., 2000). Currently, these two types of recurrence are defined solely based on their timing. Early recurrent tumors (<2 years after treatment) are usually considered true recurrences while those recurring late (>2 years post-treatment) are clinically recognized as *de novo* cancers (Imamura et al., 2003; Portolani et al., 2006; Tabrizian et al., 2015). Distinguishing these types of recurrences based on their biological characteristics remains critical because their molecular features, clinical management and prognosis can vary substantially.

Indeed, prior studies suggested that discrimination of truly recurrent versus *de novo* HCC by evolutionary trajectories and genomic heterogeneity might be more reliable than the clinical definition (Ding et al., 2019; Chen et al., 2018). A recent study revealed the differences in the immune ecosystems of primary tumors versus early relapsed HCC (true recurrence) using single-cell RNA sequencing (scRNA-seq) (Sun et al., 2021). ScRNA-seq is a powerful tool to systematically investigate the various cellular components and their interactions in the TIME (Zhang et al., 2021; Liu et al., 2022; Zhang et al., 2019). However, the TIME of *de novo* recurrent HCCs have not been studied and thus, the individual characteristics of TIME in *de novo* versus truly recurrent HCCs remain unknown. Dissecting the immune ecosystems of the two types of HCC recurrence could provide novel insights into the heterogeneity of immune evasion mechanisms between them, which might be vital to design specific and effective immunotherapeutic strategies for truly and *de novo* recurrent HCCa.

Here, we characterized the genomic profiles and TIME of two different types of recurrent HCCs with integrative multi-“omic” analyses including whole-exome sequencing (WES), scRNA-seq, bulk RNA-seq, multiplexed immunofluorescence (IF), and flow cytometry (**Figure 1A, 1B**, **Table S1**). Our study revealed the specific differences between the TIMEs in true versus *de novo* recurrence, providing guidance for future immunotherapy strategies for the management of these two different types of HCC recurrence.

**Figure 1.**
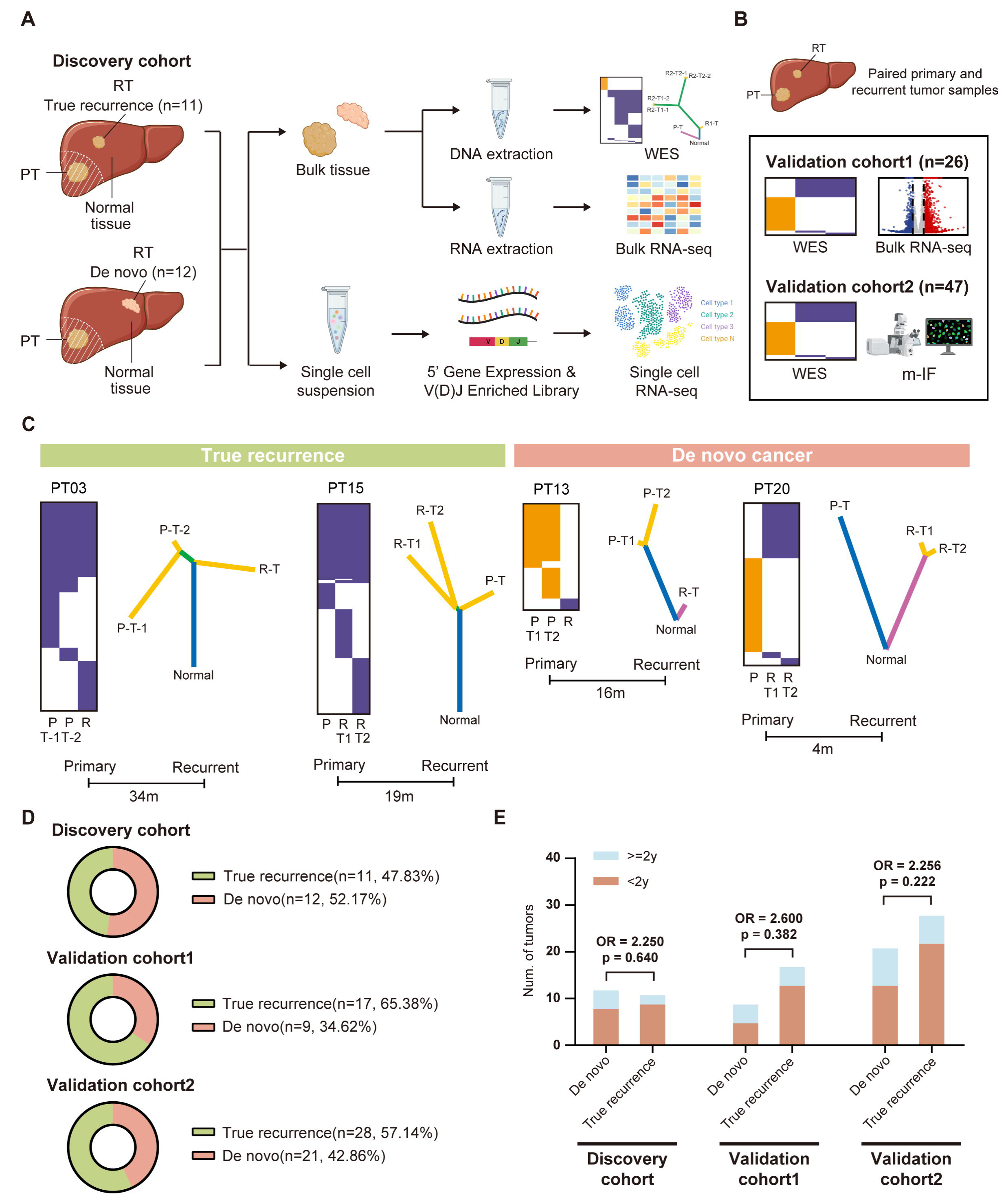
Study Workflow and Genomic Analysis Identified *De Novo* HCCs versus True Recurrences with Distinct Mutational Profiles and Evolutionary Trajectories. (A) Experimental plan for the analyses conducted in recurrent HCC samples in the discovery cohort (*n*=20 patients with 23 tumor samples). (B) Experimental plan for the analyses conducted in the validation cohort 1 and 2. (C) Heat maps of non-silent mutations (left), phylogenetic trees (right) and clinical courses (bottom) for 2 representative patients with *de novo* HCC and 2 representative patients with true recurrence. Presence (orange or purple) or absence (white) of a non-silent mutation was indicated for each tumor. Driver somatic mutations were mapped onto the phylogenetic trees. (D) Distribution of the two types of recurrence in the discovery cohort and two validation cohorts. (E) The type of recurrence was not associated with the clinically used time to recurrence (<2 years) in the discovery cohort (*p*=0.640) or in the validation cohorts 1 (*p*=0.382) or 2 (*p*=0.222).

## Results

### Distinct Mutational Profiles and Phylogenesis of Two Types of Recurrence Patterns

To determine the type of HCC recurrence, we performed WES (median depth 372×) using tumor and NAT samples in the discovery cohort (**Table S2**). Based on the mutation analysis and phylogenetic analysis between the paired primary and recurrent tumor samples, we unambiguously identified two types of recurrence: *de novo* recurrence (*n*=12, 52.2%) and true recurrence (*n*=11, 47.8%). In non-silent somatic mutations, *de novo* recurrent HCC lesions had differential mutation profiles compared to the paired primary lesions (**Figure 1C**, **Figure S1**). In contrast, the mutations largely overlapped with those in the primary tumors in the truly recurrent tumors (median of 35.8% overlap) (**Figure 1C**, **Figure S1**).

Moreover, we constructed phylogenetic trees to resolve clonal composition and hierarchy in paired tumor samples (Kumar et al., 2016; Dang et al., 2017; Suzuki et al., 2006). For *de novo* recurrence, no shared phylogenetic clades were detected for paired primary and recurrent tumors, further confirming the independent origin of the recurrent tumors. For truly recurrent HCC, paired primary and recurrent tumors were phylogenetically similar and shared common clades (**Figure 1C**, **Figure S1**). Using the same criteria for the discovery cohort, we identified 9 (34.6%) *de novo* recurrences and 17 (65.4%) true recurrences in the validation cohort 1, and 20 (42.6%) *de novo* recurrences and 27 (57.4%) true recurrences in the validation cohort 2 (**Figure 1D**, **Table S3**). The baseline characteristics of patients with *de novo* recurrences and truly recurrences were comparable in all the three cohorts (**Table S4**).

### *De Novo* Recurrences Can Often Occur Earlier Than 2 Years

In clinical practice, *de novo* HCC is defined as a recurrence more than 2 years after resection.

However, in our discovery cohort, 8 of the 12 *de novo* recurrent HCCs (66.7%) developed within 2 years (**Figure 1E**, **Figure S1**). This finding was confirmed in both two validation cohorts, with 4 (44.4%) *de novo* recurrence in the validation cohort 1 and 11 (55.0%) in the validation cohort 2 developing within 2 years of resection (**Figure 1E**). Moreover, Nei’s genetic distance was not significantly different between groups of early versus late *de novo* recurrence (*p*>0.05) (**Figure S2**). These data demonstrate that recurrence within 2 years may arise from primary cancer cells disseminating to the liver parenchyma but could also be *de novo* tumors in a substantial fraction of cases.

### True Recurrences May Occur When Primary HCCs are Undetectable

Given the finding that the truly recurrent phylogenetic lineage diverged simultaneously with the primary tumor lineage (**Figure 1**), implying the possibility of early dissemination, we constructed a model to trace the dynamics of HCC intrahepatic metastasis. First, we performed a cancer cell fraction (CCF) analysis of mutations between primary and recurrent tumors to identify a metastatic seeding pattern.

Results showed that the 12 pairs of primary and *de novo* recurrent HCC in the discovery cohort shared no sSNVs (**Table S5**). For the 11 pairs of primary and truly recurrent HCC, CCF analysis demonstrated two types of clonal metastasis in the discovery cohort (**Figure 2A**, **Table S5**). One consisted of monoclonal metastasis in the recurrent tumors of PT01, PT09, PT11, PT14, PT15, P and T16, which had multiple private sSNVs with higher CCF values and lacked shared sSNVs with lower CCF values in primary tumors (**Figure 2A**, **Table S5**), as previously reported (Hu et al., 2019). The other 5 cases were polyclonal metastases, in which recurrent tumors shared multiple sSNVs with lower CCF values in the primary tumors but lacked private mutations with higher CCF values (**Figure 2A**, **Table S5**).

**Figure 2.**
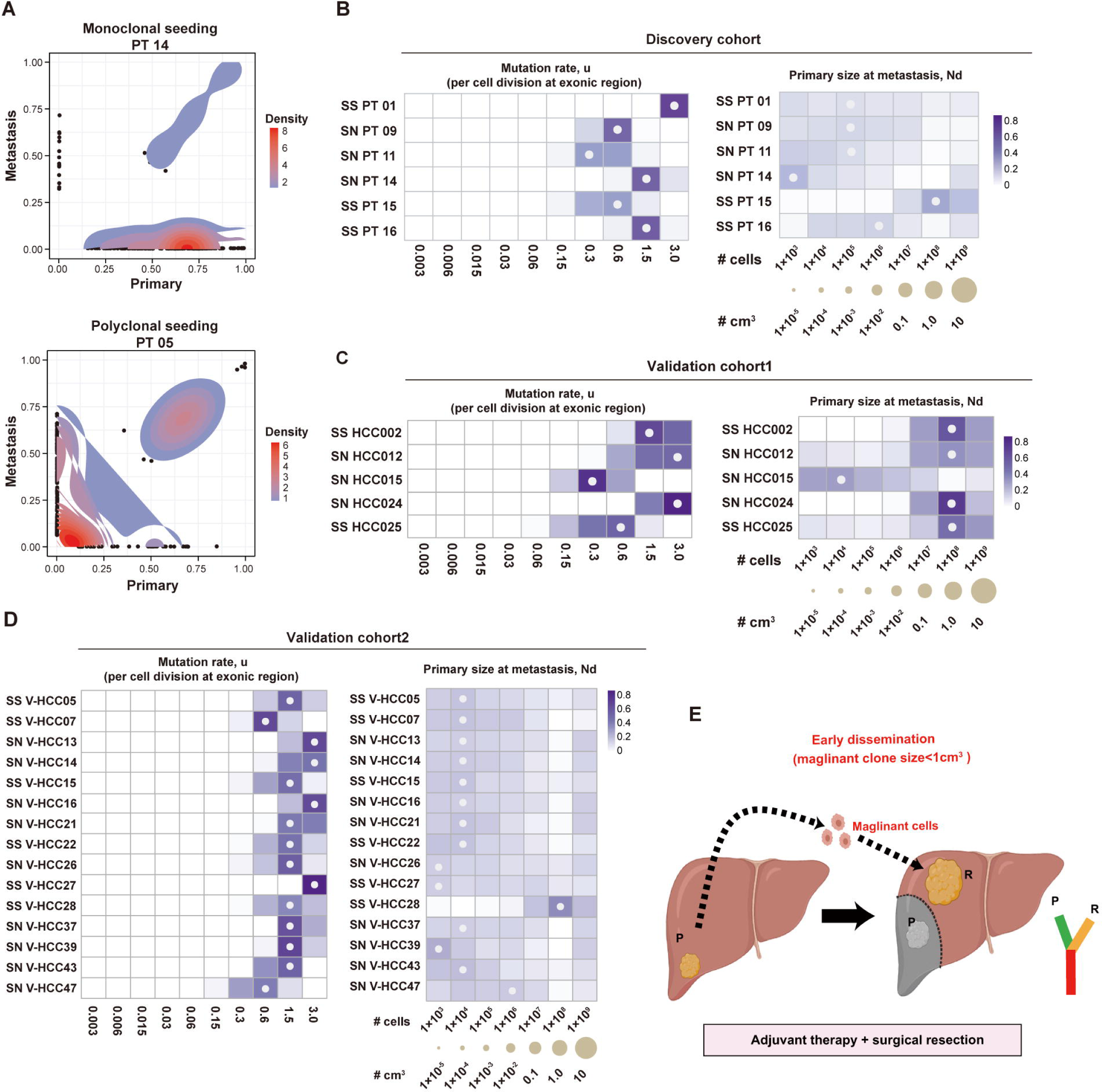
Inference of the Timing of Metastasis in HCC Patients with Monoclonal Seeding of Primary Tumors. (A) Representative density plots of cancer cell fraction (CCF) of somatic single nucleotide variant (sSNV) estimated for paired primary and recurrent tumors with monoclonal seeding pattern (PT14) and polyclonal seeding pattern (PT05) in the discovery cohort. For monoclonal seeding pattern, recurrent tumors had multiple private clonal sSNVs and lacked shared subclonal sSNVs with primary tumors while for polyclonal seeding pattern, recurrent tumors shared multiple subclonal sSNVs with primary tumors but lacked private clonal mutations. Other density plots were in Supplementary Figure 3. (B) Heatmap of the inferred posterior probability distributions for the mutation rate u (per cell division in exonic regions) and Nd (the tumor size of primary tumor when metastasis occurred) in patients PT01, PT09, PT11, PT14, PT15and PT16 in the discovery cohort. Primary tumors began to disseminate when they reached 1x10^3^–1x10^8^ cells (0.00001–1 cm^3^ in diameter). The median of the posterior distribution is marked with a white circle at the corresponding value. The tumor evolution scenario in each primary and recurrent pair was determined by SCIMET. The corresponding tumor volumes of each Nd is calculated and marked below the heatmap. (C-D) Heatmap of the inferred posterior probability distributions for the mutation rate u (per cell division in exonic regions) and Nd (the tumor size of primary tumor when metastasis occurred) in patients with monoclonal seeding pattern from patients with truly recurrent HCC in the validation cohort 1 (C) and 2 (D). Primary tumors began to disseminate when they were less than 1×10^8^ cells (<1 cm^3^ in diameter). (E) Schematic diagram of early dissemination of primary HCC in the liver parenchyma when they are <1cm in diameter that could result in the true HCC recurrence.

SCIMET model was applied to infer further the timing of metastasis in our paired samples (Hu et al., 2019). The SCIMET algorithm assumed that metastasis was originally seeded by a single cell; thus, only PT01, PT09, PT11, PT14, PT15, and PT16 in the discovery cohort, which showed apparent monoclonal seeding patterns, were included in this analysis. The evolutionary mode inferred from the SCIMET model demonstrated that the recurrent tumors of PT01, PT15, and PT16 underwent stringent subclonal selection; in contrast, the recurrent tumors of PT09, PT11, and PT14 tended to develop under a neutral scenario (**Figure 2B**). SCIMET model showed a relatively high patient-specific mutation rate (u) in these patients. A rate of 1.5 mutations per cell division is expected in exonic regions during tumor progression. Based on the number of somatic mutations seen in the paired tumor samples, we further quantified the size of the primary tumor when cancer cells began to spread to other areas (Nd) in the liver. This time-point was defined as the time of dissemination. Notably, in these patients, early dissemination occurred when primary tumors were less than 1 cm^3^ in volume (1×10^8^ cells), a size at which HCCs are rarely diagnosed clinically. Moreover, 3 of the 6 cases were intrahepatic recurrences (PT09, PT14, and PT15), implying early distant dissemination from the primary tumor. We confirmed this finding in the two validation cohorts, in which we found that 5/17 (29.4%) true recurrences in the validation cohort 1 and 15/27 (55.6%) true recurrences in the validation cohort 2 were monoclonal metastases (**Figure 2C, 2D**, **Table S5**). In all cases, early dissemination was estimated to have developed when the primary tumor was less than 1 cm^3^ in volume, consistent with the results in the discovery cohort. Altogether, these data showed that early seeding from the primary HCCs, giving rise to rapid recurrences even in the case of complete surgical removal of primary tumors (**Figure 2E**).

### Profiling by scRNA-seq of TIMEs in the Two Types of Recurrent HCC

To compare the TIMEs of truly and *de novo* recurrent HCCs, we performed 5’and VDJ scRNA-seq analysis in 23 recurrent tumor samples and 11 NAT samples. After quality control and filtering, we obtained 317,558 high-quality cells (median: 9,074 cells [range; 1,712-15,272 cells]/sample) with 1,648 detected genes on average (**Table S6**). Then, we performed cell classification and marker gene identification using Seurat 4.0 and detected and visualized 4 major clusters using Umap (Uniform Manifold Approximation and Projection for Dimension Reduction) (**Figure 3A**). To generate a global cell atlas in recurrent HCCs, we further sub-clustered each major cell compartment and revealed 43 cell subtypes (**Figure 3B**, **Table S7**). As demonstrated in **Figure 3A**, non-immune cells included hepatic stellate cells (HSCs; identified by expression of *ACTA2* and *PDGFRB*), endothelial cells (ECs; identified by expression of *PECAM1* and *CDH5*), and malignant cells (identified by expression of *ALB*, *TTR* and *APOA2*). As expected, malignant cells accounted for most cells of the non-immune cells, while HSCs and ECs only accounted for a smaller proportion (28.33%) (**Figure 3A**, **Figure S3A**). Tumor origin was confirmed by detecting copy number variations (CNVs), inferred from the scRNA-seq data (**Figure S3B**). On the other hand, the immune cells included T cell/innate lymphoid cells (ILCs) (identified by expression of *CD3D/E*, *TYROBP*), myeloid cells (identified by expression of CD68 and *LYZ*), neutrophils (identified by expression of CSF3R), plasmacytoid dendritic cells (pDCs) (identified by expression of *LILRA*4 and *IL3RA*), B cells (identified by expression of *MS4A1* and *CD79A*), plasma cells (identified by expression of *MZB1* and immunoglobins) and mast cell (identified by expression of *CPA3*) (**Figure 3A, 3B**). These immune cell subtypes were present in all truly and *de novo* recurrent HCC samples (**Figure 3C**, **Figure S3A**). Overall, the two types of recurrent HCCs shared similar proportions of lymphoid-derived, myeloid-derived and B-lineage cells (**Figure S3C**).

**Figure 3.**
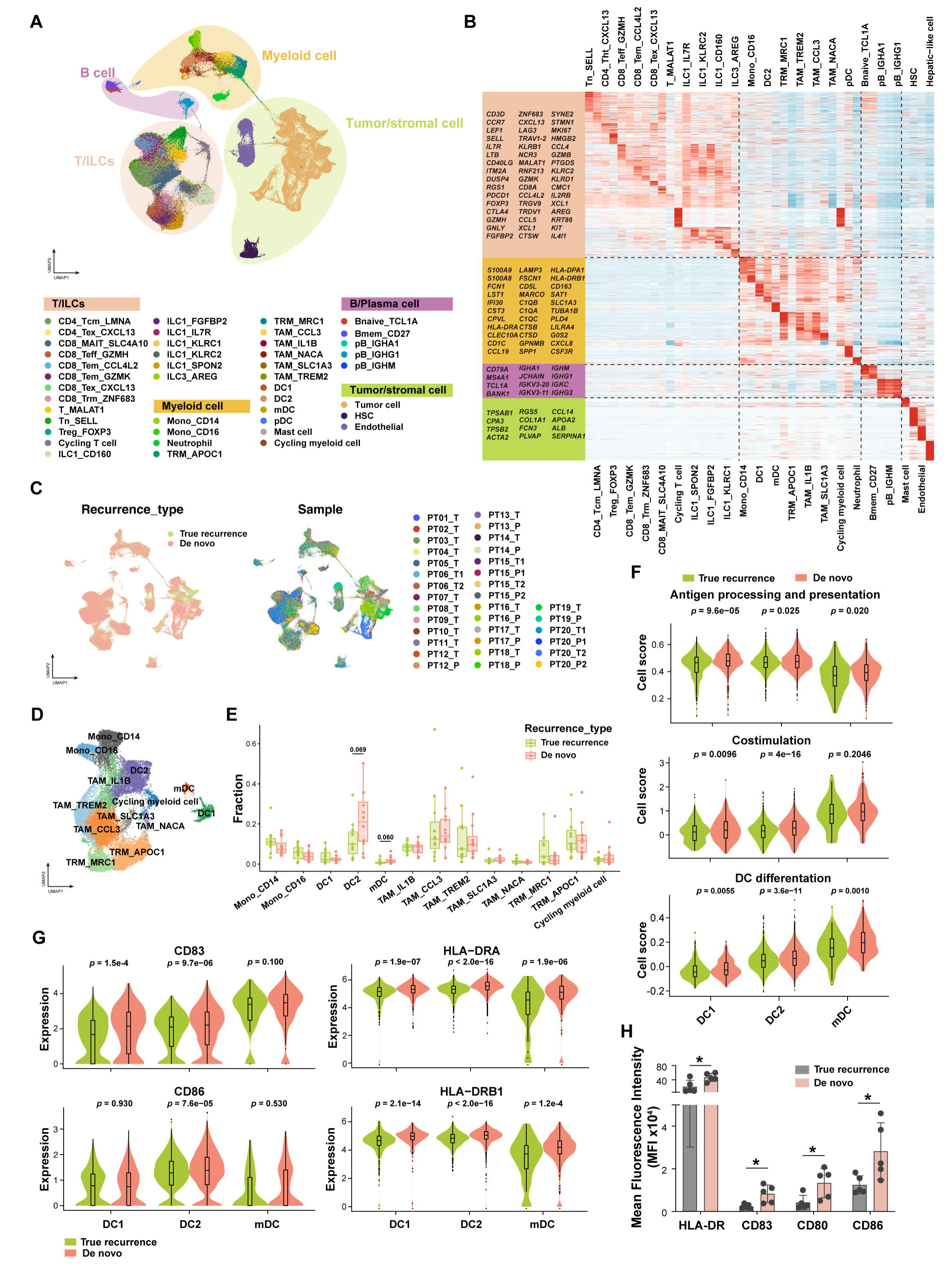
ScRNA-seq Landscape of Recurrent HCC Immune Microenvironment. (A) Uniform Manifold Approximation and Projection (UMAP) plot of cells color-coded for the indicated cell types in the microenvironment of recurrent HCC, displaying 10 major cell types and 43 subtypes in total. (B) Heatmap showing the expression patterns of signature genes in the indicated cell types. (C) UMAP plot showing cell origins by color, the origin of recurrent types (left panel) and the origin of patient samples (right panel). (D) UMAP plot showing the subtypes and corresponding annotations of myeloid cells in recurrent HCC patients. (E) Comparison of the cell fractions of myeloid subtypes between true recurrent tumor samples (green) and *de novo* recurrent tumor samples (red) by Wilcoxon test. (F) Violin plots showing the expression scores of antigen processing and presentation signature, co-stimulation signature and differentiation signature of DC1, DC2 and mDC in true recurrent tumor samples (green) and *de novo* recurrent tumor samples (red). Comparisons were performed by Wilcoxon test with p values indicated above. (G) Violin plots showing the expression of co-stimulatory and MHC-II molecular genes in three types of DCs in true recurrent tumor samples (green) and *de novo* recurrent tumor samples (red). Comparisons were performed by Wilcoxon test with p values indicated above. (H) Flow cytometry analysis confirmed the increased expressions of HLA-DR, CD83, CD80 and CD86 molecules in DCs for *de novo* recurrent tumor samples (pink) when compared with truly recurrent tumor samples (gray). * indicates a p<0.05 from Student’s t test.

### Distinct DC Frequencies in the TIME of Truly Versus De Novo Recurrent HCCs

We next performed unsupervised clustering of the myeloid cells and T/ILC cell populations. Myeloid cells could be re-clustered into 13 clusters, including 5 clusters of tumor-associated macrophages (TAMs) (TAM_CCL3, TAM_IL1B, TAM_NACA, TAM_SLC1A3, TAM_TREM2), 3 clusters of dendritic cells (DCs) (DC1, DC2, mDC), 2 clusters of tissue-resident macrophages (TRMs) (TRM_APOC1, TRM_MRC1), 2 clusters of monocytes (Mono_CD14, Mono_CD16) and 1 cluster of cycling cells (**Figure 3D, Figure S3D, Table S7**). Macrophages were significantly enriched in tumor tissues compared to the paired NATs in both recurrence types and showed similar proportions among immune fractions in truly and *de novo* recurrent tumor samples (**Figure 3E, Figure S3E**). TRMs were also similar in truly and *de novo* recurrent tumor samples (**Figure 3E**, **Figure S3E**). Among all myeloid cell subtypes, only DC2 and mDC had marginally higher proportions in *de novo* recurrent tumors than in truly recurrent tumors, while DC1 had similar frequencies between these two types of recurrent tumors. Flow cytometry showed a significantly increased fraction of mDCs (CD45^+^CD14^-^CD11c^+^CCR7^+^) in *de novo* recurrent tumors versus truly recurrent tumors and a tendency for an increased fraction of DC1 (CD45^+^CD14^-^CD11c^+^XCR1^+^) and DC2 (CD45^+^CD14^-^CD11c^+^CD1c^+^) in *de novo* recurrent tumors (**Figure S3F**). Interestingly, we found that all the DC subtypes (DC1, DC2, and mDC) demonstrated increased antigen processing and presentation, co-stimulation, and differentiation transcriptional signatures in *de novo* versus truly recurrent tumors (**Figure 3F**). Consistently, the expression levels of classical co-stimulatory and MHC-II molecules of DC such as *CD83*, *CD86*, *HLA-DRA,* and *HLA-DRB1* were also higher for *de novo* recurrent tumors (**Figure 3G**). We validated these scRNA-seq data by using flow cytometry analysis (**Figure 3H**). Collectively, these data show largely similar myeloid cell profiles except for DCs, whose frequency and function were reduced in truly recurrent HCC.

### Distinct Lymphocyte Subset Frequencies in the TIME of Truly Versus De Novo Recurrent HCCs

ScRNA-seq analysis of the lymphocyte subsets identified 19 clusters, including 6 of CD8^+^ T cells, 7 of ILCs, 2 of CD4^+^ T cells, and 1 cluster of regulatory T cell (Treg), naïve T cell (Tn), T_MALAT1 cell and cycling T cells (**Figure 4A**, **Table S7**). These clusters were present in all truly and *de novo* recurrent tumor samples (**Figure 4B**). CD8^+^ T cells were more abundant in NATs, while CD4^+^ T cells and Treg cells were enriched in tumors of both recurrence types (**Figure S4A**), consistent with previous reports (Zheng et al., 2017). However, when we compared the total fractions of T cell subtypes between the two types of HCC recurrence, we found that the proportion of CD8^+^ T cells was higher in truly recurrent tumors than in *de novo* recurrent HCCs, and that the fraction of CD4^+^ T cells showed the opposite trend (**Figure 4C**). We validated this finding by multiplexed IF suing samples from validation cohort 2 (**Figure 4D**). The fractions of other T cell subtypes in tumor samples did not significantly differ between the two recurrence types (**Figure S4B**).

**Figure 4.**
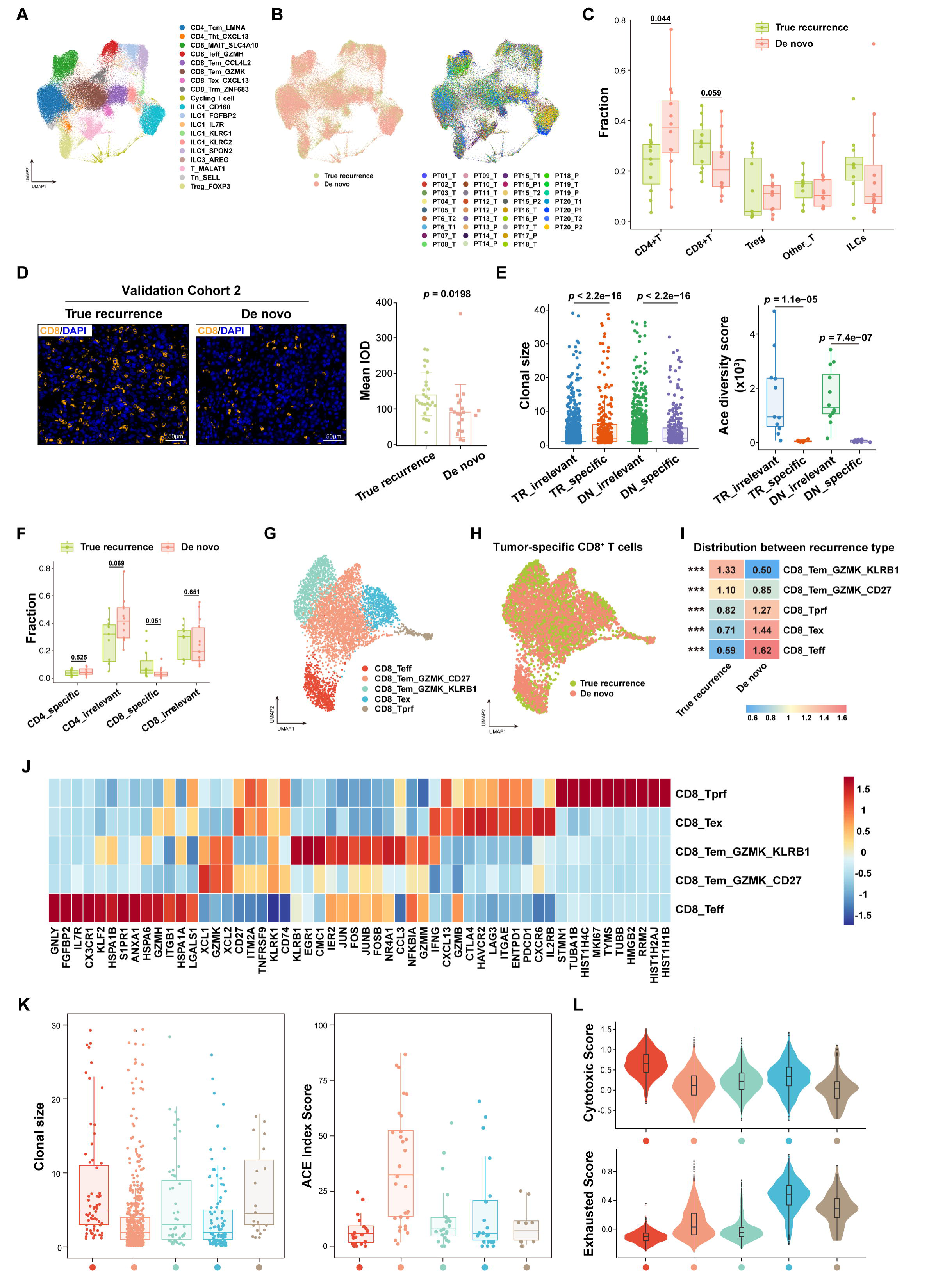
Identification and Characteristics of Tumor-specific CD8^+^ T cell Subtypes in Recurrent HCC. (A) UMAP plot showing the subtypes and corresponding annotations of T cells/innate lymphoid cells (ILCs) in recurrent HCC patients. (B) UMAP plot showing cell origins of T cell/ILCs by color, the origin of recurrent types (left panel) and the origin of patient samples (right panel). (C) Comparison of the cell fractions of CD4^+^ T, CD8^+^ T, regulatory T (Treg), ILCs and other T cell subtypes between true recurrent tumor samples (green) and *de novo* recurrent tumor samples (red) by Wilcoxon test. (D) Representative multiplexed immunofluorescent images for CD8^+^ T cells in true recurrent tumor area (left upper panel) and *de novo* recurrent tumor area (left bottom panel). Boxplot (right panel) showing the density of CD8^+^T cells in the validation cohort 1 (*n*=26 patients), and comparison was performed using Student’s t test. (E) The clone size (upper panel) and Ace diversity score (lower panel) of tumor-specific and tumor-irrelevant CD4^+^/CD8^+^ T cells in tumor samples of two recurrent types. *p* values were calculated by Wilcoxon test. (F) Comparison of the cell fractions of tumor-specific CD4^+^ T cells, tumor-irrelevant CD4^+^ T cells, tumor-specific CD8^+^ T cells and tumor-irrelevant CD8^+^T cells between true recurrent tumor samples (green) and *de novo* recurrent tumor samples (red) by Wilcoxon test. (G) UMAP plot showing the five subtypes and corresponding annotations of tumor-specific CD8^+^T cells in recurrent HCC patients. Red color coded for CD8_Teff, orange coded for CD8_Tem_GZMK_CD27, green coded for CD8_Tem_GZMK_KLRB1, blue coded for CD8_Tex and brown coded for CD8_Tprf. (H) UMAP plot showing the distribution of tumor-specific CD8^+^ T cells in true and *de novo* recurrent tumors. (I) Heatmap showing the relative enrichment of five subtypes of tumor-specific CD8^+^T cells in true recurrent and *de novo* recurrent tumors (*** indicates a *p*<0.001). Red color represents the enrichment while blue color represents the opposite. (J) Heatmap showing mean relative expression of certain signature genes in the five cell subtypes identified in panel G. (K) The clone size (left panel) and Ace diversity score (right panel) of five cell subtypes identified in panel G. (L) Violin plots showing the expression of cytotoxic (upper panel) and exhaustion scores (lower panel) of the five cell subtypes identified in panel G.

Emerging evidence suggests that tumor-specific T cells are the critical T cell component in anti-tumor immunity (Liu et al., 2022; Simoni et al., 2018; Cohen et al., 2022; van der Leun et al., 2020). Thus, we identified and characterized the tumor-specific T cells by applying a TCR-based selection strategy based on two assumptions from published literature (Liu et al., 2022; Cohen et al., 2022; Duhen et al., 2018): **1**) cells sharing TCRs with exhausted T cells; and **2**) cells highly expressing the tumor-specific T cell signature gene, including *ENTPD1*, *ITGAE*, *PDCD1*, *CXCL13*, *MAF,* and *ZBED2*. Based on this strategy, we classified T cells into tumor-specific and tumor-irrelevant T cells and found that the clone size of tumor-specific CD4^+^ T/CD8^+^ T cells was significantly larger than that of tumor-irrelevant CD4^+^ T/CD8^+^ T cells (**Figure 4E**). In addition, clone diversity of tumor-specific T cells was lower than that of tumor-irrelevant T cells (**Figure 4E**), consistent with previous reports (Liu et al., 2022; Simoni et al., 2018; Cohen et al., 2022; van der Leun et al., 2020). Interestingly, by comparing the tumor-specific and tumor-irrelevant T cells between the two recurrence types, we found an increased abundance of tumor-specific CD8^+^ T cells in truly recurrent versus *de novo* recurrent HCCs (**Figure 4F, Figure S4C**). This finding was further supported by the higher expression of tumor specific CD8^+^ T cell signature genes in truly recurrent tumors from validation cohort 1 (**Figure S4D**) (Simoni et al., 2018). In contrast, we found an increased fraction of tumor-irrelevant CD4^+^ T cells in *de novo* recurrent tumors (**Figure 4F**).

### Distinct Tumor-Specific CD8^+^ T Cell Subtypes in the TIME of Truly Versus De Novo Recurrent HCCs

We subsequently focused on defining the tumor-specific CD8^+^ T cells given the findings above. By performing re-clustering analyses of sc-RNAseq data, we detected 5 tumor-specific CD8^+^ T cell subtypes: CD8_Teff, CD8_Tex, proliferative CD8^+^ T cells (CD8_Tprf), CD8_Tem_GZMK_KLRB1 and CD8_Tem_GZMK_CD27 (**Figure 4G, Figure S4E**). CD8_Tem_GZMK_KLRB1 and CD8_Tem_GZMK_CD27 cells were increased in truly recurrent tumors, whereas CD8_Tex and CD8_Teff cells were enriched in *de novo* recurrent HCCs (all *p*<0.001; **Figure 4H, I**). CD8_Teff cells showed elevated expression levels of cytotoxic genes, such as *GNLY*, *FGFBP2*, *CX3CR1,* and *GZMH*, while CD8_Tex cells showed high expression of T cell exhaustion markers, such as *PDCD1*, *CXCL13, CTLA4, HAVCR2,* and *LAG3* (**Figure 4J**). In addition, we detected two subtypes of memory CD8^+^ T cells, both expressing *GZMK* and chemokines, including *XCL1* and *CCL4L2*. CD8_Tem_GZMK_CD27 cells displayed high expression levels of effector memory-associated coactivated genes *CD27*, *TNFRSF9,* and *CD74* and the relatively low expression of checkpoint genes (**Figure 4J**). CD8_Tem_GZMK_KLRB1 cells demonstrated the highest expression of an inhibitory receptor *KLRB1* (*CD161*), activator protein 1 genes (*JUN*, *JUNB*, *FOS*, *FOSB*), *CMC1,* and *NR4A1*, suggesting a different memory-like T cell phenotype (Ruiz et al., 2014) (**Figure 4J**). In addition to the marker expression, we next analyzed the clonal diversity and clone size of these tumor-specific CD8^+^ T cell subtypes and found that the two CD8_Tem cells also showed distinct features (**Figure 4K**). CD8_Tem_GZMK_CD27 had the highest clonal diversity, implying an intermediate cell state, whereas CD8_Tem_GZMK_KLRB1 (also named CD161^+^CD8^+^ T cell) displayed a low clonal diversity and a moderate degree of clonal expansion between CD8_Tex and CD8_Teff. Functionally, these two types of CD8_Tem cells exhibited low expression levels of both exhausted and cytotoxic signatures (Sun et al., 2021; Wu et al., 2021) (**Figure 4L**). More importantly, the expression level of the cytotoxic signature was even lower than that of CD8_Tex, implying the weak cytotoxicity of these cells (**Figure 4L**). Previous reports demonstrated the impaired cytotoxicity of CD8^+^CD161^+^ T cells and their immunosuppressive function of inhibiting the effector T cells infiltrated into tumors (Sun et al., 2021; Mathewson et al., 2021; Di et al., 2022). Thus, we further examined the CD161-associated transcriptional signature and found considerable similarities in gene expression with the CD161^+^CD8^+^ T cell subsets in recurrent HCC tissue (**Figure S4F**), consistent with the weak cytotoxicity of CD161^+^CD8^+^ T cells. In addition, to further confirm the definitions of these five functional annotations, we traced their corresponding clusters in the overall CD8^+^ T cell annotations. All the CD8_Tprf belonged to the cycling T cells, and most CD8_Teff and CD8_Tex cells belonged to CD8_Teff_GZMH and CD8_Tex_CXCL13 cells, respectively, confirming the accuracy of the functional classification (**Figure S4G**). Most CD161^+^CD8^+^ T cells belonged to CD8_Tem_CCL4L2 cells that resided between cytotoxic and exhausted states (**Figure S4H**). Altogether, these data identify CD8_Teff, CD8_Tex, CD8_Tprf, CD161^+^CD8^+^ T cell, and CD8_Tem_GZMK_CD27 as five transcriptionally distinct tumor-specific CD8^+^ T cells in recurrent HCC, and indicate that CD161^+^CD8^+^ T cells might be in the memory state with weak cytotoxicity and immunosuppressive phenotype. In addition, they highlight other differences in the TIMEs of truly versus *de novo* recurrent HCCs, with decreased abundance of CD8^+^ T cells in *de novo* recurrent tumors.

### Tumor-specific CD8^+^ T cells Reside in a Cytotoxic but Exhausted State in De Novo Recurrent HCC and in a Low Cytotoxic State in Truly Recurrent HCC

In light of the TIME differences between these two recurrent types, we next examined the transcriptional states and dynamic cell transitions of tumor-specific CD8^+^ T cells. First, differentially expressed gene (DEG) analysis showed that CD8^+^ T cells in *de novo* recurrent HCC upregulated *CTLA4*, *CXCL13*, *PDCD1*, *HAVCR2*, *GZMB,* and *GNLY*, suggesting an exhausted cytotoxic T cell phenotype (**Figure 5A, 5B**). In contrast, CD8^+^ T cells in truly recurrent HCCs showed upregulation in *XCL1, EOMES, ID3, CMC1*, *CD74,* and *KLRB1*, indicating a suppressive memory T cell phenotype (Zhang et al., 2021; Ruiz et al., 2014; Istaces et al., 2019; Dahmani et al., 2019) (**Figure 5A, 5B**). Notably, *KLRB1* was specifically upregulated in truly recurrent HCCs (**Figure 5C**). Then, to further compare the transcriptional functional states of CD8^+^ T cells among the two recurrence types, we evaluated the expression scores in cytotoxic and exhausted signatures (Sun et al., 2021; Wu et al., 2021). In concordance with the result of the DEG analysis, we found an increased expression in exhausted signature and a slight increase in cytotoxic signature in CD8^+^ T cells in *de novo* versus truly recurrent samples (**Figure 5D**).

**Figure 5.**
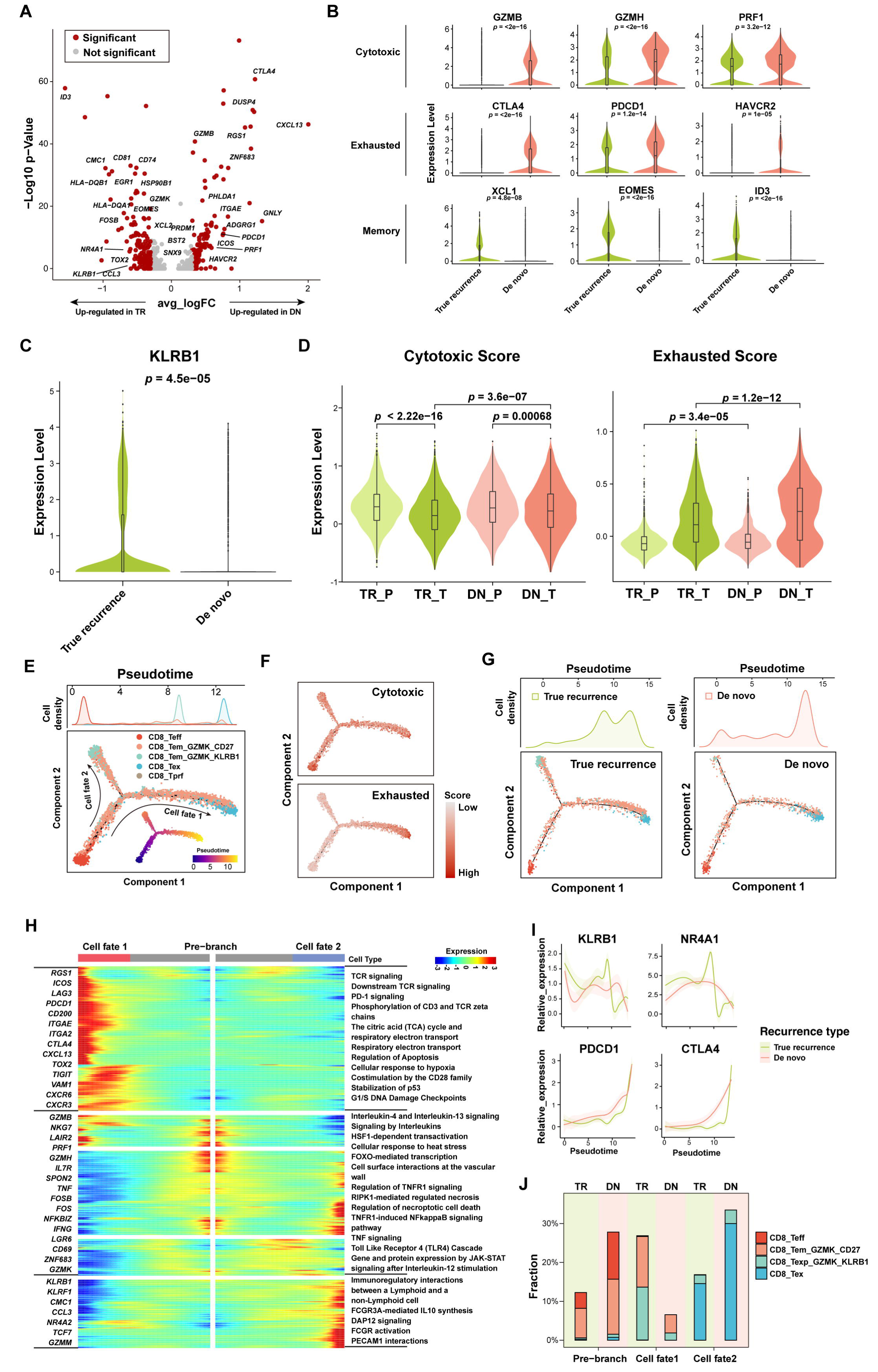
Tumor-specific CD8^+^ T cells Reside in Cytotoxic/exhausted States for *De Novo* Recurrent HCC while Reside in Low Cytotoxic State for Truly Recurrent HCC. (A) Volcano plot showing the differentially expressed genes of tumor-specific CD8^+^ T cells between truly recurrent and *de novo* recurrent tumors. The most significant genes were indicated in the plot. (B) Comparison of the expression level of cytotoxic, exhausted and memory-associated genes between truly recurrent tumors (green) and *de novo* recurrent tumors (red) with Wilcoxon test. (C) Violin plot showing the expression level of *KLRB1* in truly recurrent (green) and *de novo* recurrent tumors (red). (D) Violin plot showing the expression score of cytotoxic and exhausted signatures in tumor samples and non-tumor adjacent liver tissue samples of true recurrence and *de novo* recurrence. Comparison was performed by Wilcoxon test. (E) Pseudotime trajectory for tumor-specific CD8^+^ T cells in recurrent HCC samples based on Monocle (Bottom), with cell density plot of five subtypes of tumor-specific CD8^+^T cells along with the pseudo-time (Top). (F) Pseudotime plot showing the expression of cytotoxic (upper panel) and exhausted signatures (lower panel) for tumor-specific CD8^+^ T cells along the trajectory. (G) Pseudotime plot of tumor-specific CD8^+^ T cells in truly recurrent tumors (upper panel) and *de novo* recurrent tumors (lower panel) respectively, with cell density plot showing the distribution of five subtypes of tumor-specific CD8^+^ T cells along with the pseudo-time (Top). (H) Gene expression dynamics along the trajectory of tumor-specific CD8^+^ T cells outlined in E, from the pre-branch phase to cell fate 1 and cell fate 2. Genes cluster into three gene sets, each characterized by specific expression profiles, as depicted by some marker genes (left) and enriched pathways (right) characteristic for each cluster. (I) *KLRB1*, *NR4A2*, *PDCD1* and *CTLA4* expression levels associated with CD8^+^ T cell transition along the pseudotime in truly recurrent tumors (green) and *de novo* recurrent tumors (red). (J) Histogram showing the distribution of four subtypes of tumor-specific CD8^+^ T cells in three phases identified in H, for truly recurrent and *de novo* recurrent tumors.

To further confirm the distinct immune states of tumor-specific CD8^+^ T cells in the two recurrent HCC types, we inferred the state trajectories using Monocle to investigate the dynamic cell transitions. Due to the absence of naïve cells among tumor-specific CD8^+^ T cells, pseudotime analysis revealed that the CD8_Teff cells were at the beginning of the trajectory path, while the CD8_Tex cells and CD8_Tem_GZMK_CD161 cells were at the end of the trajectory path (**Figure 5E**). Based on TCR clonality (**Figure S4I**) and trajectory data, the transition was confirmed to initiate with CD8_Teff cells, through an intermediate effector memory state featured by CD8_Tem_GZMK_CD27 cells, and finally to reach two terminal states, one being the exhausted state featured by CD8_Tex cells and the other the memory state featured by CD8_Tem_GZMK_CD161 cells (**Figure 5E**). During this transition, the cytotoxic signature score was reduced while the exhausted signature score increased (**Figure 5F**). Notably, CD8_Tem_GZMK_CD161 cells expressed even lower cytotoxic scores than CD8_Tex cells, demonstrating their relatively quiescent state with weak cytotoxicity (**Figure 4L and 5F**).

### Differential Gene Expression Patterns in Tumor-Specific CD8^+^ T Cells Based on Activation State and Type of HCC Recurrence

Given these findings, we next investigated the trajectories of CD8^+^ T cells in samples from truly and *de novo* recurrent HCCs to analyze the transition cell states. Interestingly, cytotoxic CD8^+^ T cells were predominantly found in *de novo* recurrent tumors with few cells at the terminal end of CD8_Tem_GZMK_CD161, whereas CD8_Tem_GZMK_CD161 cells were primarily detected in truly recurrent HCCs (*p*<2.2×10^-16^, ks-test) (**Figure 5G**). Our analysis showed that the intermediate CD8_Tem_GZMK_CD27 cells were more likely to transit to exhausted CD8^+^ T cell states in *de novo* recurrent HCC samples. In contrast, the intermediate CD8_Tem_GZMK_CD27 cells in truly recurrent tumor samples showed two different paths, one transitioning to the exhausted state and the other to the memory state with weak cytotoxicity (**Figure 5G**). We then examined the transcriptional changes associated with different transitional CD8^+^ T cell states (**Figure 5H**). Based on the branching point of the two trajectory paths described above, CD8^+^ T cells were categorized into three phases: pre-branch phase, fate 1 phase (the path with terminal end of CD8_Tex), and fate 2 phase (the path with terminal end of CD8_Tem_GZMK_CD161). The T cells in pre-branch phase presented the highest expression levels of *GZMB*, *GZMH*, *NKG7*, *PRF1,* and *IFN*, matching the phenotypes of cytotoxic and early activated T cells (**Figure 5H**). The T cells in fate 1 phase showed high levels of classical checkpoint and exhausted-related genes such as *LAG3*, *PDCD1*, *CTLA4*, *TIGIT,* and *CXCL13*. Pathway analysis revealed that TCR and PD-1 signaling pathways were enriched in fate 1 phase, consistent with the exhausted phenotype of CD8_Tex cells at this trajectory end (**Figure 5H**). Notably, CD8_Tem_GZMK_CD161 cells were mainly in fate 2 phase, and demonstrated high expression levels of *KLRB1*, *CCL3*, *TCF7*, and *NR4A2* and low expression levels of *GZMB*, *GZMH,* and *NKG7*, consistent with a memory phenotype and low cytotoxicity (Sun et al., 2021; Ruiz et al., 2014; Mathewson et al., 2021; Di et al., 2022) (**Figure 5H**). Furthermore, pathway analysis showed that Immunoregulatory and FCGR3A-mediated IL10 synthesis were upregulated in cells in the fate 2 phase, further confirming the weak cytotoxicity of cells in fate 2 phase (**Figure 5H**). IL10 is an immunosuppressive cytokine which inhibits the expression of *GZMB* and *IFN* in CD8^+^ T cells, dampening their activation and anti-tumor effector functions (Xiao et al., 2016; Wei et al., 2019).

Consistent with the above findings, the expression levels of *KLRB1* and *NR4A2* were upregulated and reached a peak in the fate 2 phase in T cells in truly recurrent samples; in contrast, these genes were not upregulated through the trajectory path of T cells in *de novo* recurrent HCCs (**Figure 5I**). Moreover, the expression levels of the checkpoint molecules *PDCD1* and *CTLA4* were gradually increased from the cytotoxic to exhausted states and were higher in the T cells in *de novo* versus truly recurrent samples (**Figure 5I**). Finally, we further compared the distribution of the CD8^+^ T cells in different phases between the two recurrence types. We found that CD8^+^ T cells in *de novo* recurrent samples predominantly resided in the pre-branch phase and exhausted fate 1 phase (**Figure 5J**). Conversely, CD8^+^ T cells in truly recurrent samples mainly resided in fate 2 phase, with some in the exhausted phase, suggesting a unique developmental program from the intermediate CD8_ Tem_GZMK_CD27 cells to a relatively quiescent memory phenotype, in addition to the traditional transition process to T cell exhaustion (**Figure 5J**).

Altogether, our analysis of CD8^+^ T cells in truly and *de novo* recurrent HCC samples demonstrated distinct immune and transcriptional states during the transition trajectories. CD8^+^ T cells in *de novo* recurrent tumor samples were characterized by cytotoxic and exhausted phenotypes, while CD8^+^ T cells in truly recurrent HCC samples were characterized by abundance of memory immunosuppressive cells (CD8_Tem_GZMK_CD161) with weak cytotoxicity.

### Distinct Patterns of Immunosuppression in the TIME of the Two Types of Recurrent HCC

To further decipher the pathways associated with the exhausted state of CD8^+^ T cells in *de novo* recurrent HCC samples, we interrogated the connections among different immune cells within the TIME. We first found that the top two immune cell subtypes positively correlated with the exhaustion of tumor-specific CD8^+^ T cells were TAM and mDC (**Figure 6A**). Ligand-receptor (L-R) analysis also showed that myeloid cells, such as TAM and mDC, interacted closely with CD8^+^ T cells via LGALS9 (Galectin9) signaling, and these interactions appeared to be stronger in *de novo* versus truly recurrent HCC samples (**Figure 6B**). Moreover, the expression levels of LGALS9 were higher in myeloid cells (i.e., mDC, TAM_TREM2, TAM_CCL3) from *de novo* than truly recurrent tumors (**Figure 6C**). Multiplexed IF also demonstrated that Galectin9^+^CD68^+^ myeloid cells were in proximity of Tim3^+^CD8^+^ T cells, supporting their crosstalk in *de novo* recurrent tumors (**Figure 6D**). We did not detect a significant interaction between myeloid cells with CD8^+^ T cell via PD-L1/PD1 when analyzing scRNA-seq data, likely due to the dynamic and low expression of PD-L1. Therefore, considering the important role of the PD-L1/PD-1 axis in advanced HCC treatment, we performed multiplexed IF and found that PD-L1^+^ myeloid cells resided near PD1^+^CD8^+^ T cells in *de novo* recurrent tumors (**Figure S5A**). Overall, these findings suggested the important role of myeloid cells via LGALS9 and PD-L1 axes in the exhaustion of CD8^+^ T cells in the TIME of *de novo* recurrent HCC.

**Figure 6.**
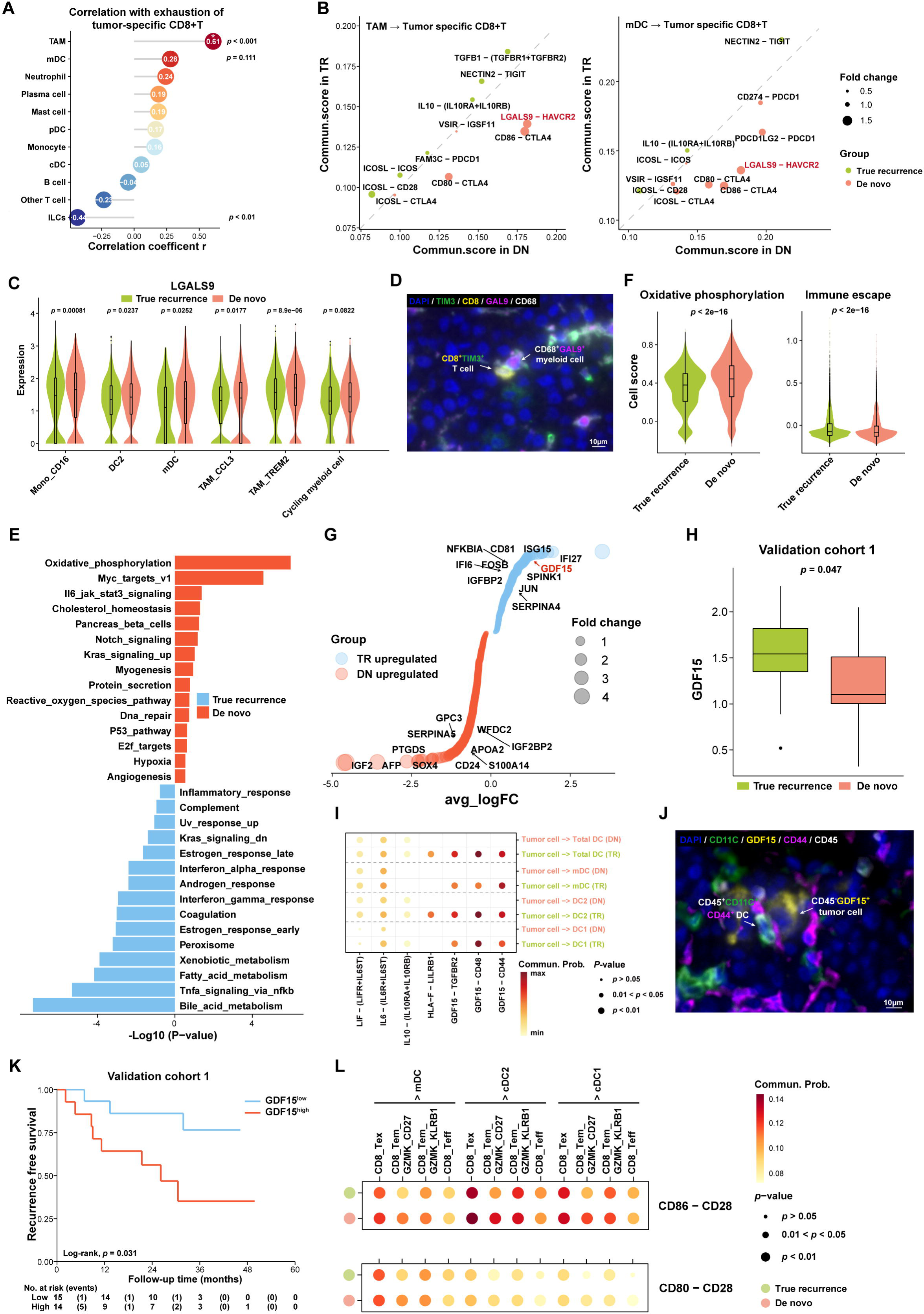
Distinct Cell-cell Interaction Patterns in *De Novo* versus Truly Recurrent HCC Microenvironments. (A) Correlations of immune cell subsets in their cellular proportions with the exhaustion score of tumor-specific CD8^+^ T cells in recurrent tumor samples. (B) Ligand and receptor (L-R) pairs between TAM and tumor-specific CD8+ T cells (left panel), and between mDC and tumor-specific CD8+ T cells (right panel) in *de novo* recurrent tumors and truly recurrent tumors. The L-R pairs below the diagonal line indicated its superiority in *de novo* recurrent tumors while those above the diagonal line indicated its superiority in truly recurrent tumors. (C) Violin plot showing the *LGALS9* expression levels in six myeloid cell subtypes for truly recurrent tumors (green) and *de novo* recurrent tumors (red). Comparison was performed by Wilcoxon test. (D) Representative multiplexed immunofluorescent images showing that LGALS9^+^CD68^+^ myeloid cells resided near HAVCR2^+^CD8^+^ T cells. (E) Pathways enriched in *de novo* recurrent versus truly recurrent malignant cells, based on gene set enrichment analysis (GSEA) using HALLMARK gene set. (F) Violin plot showing the expression score of proliferation and immune escape signatures in truly recurrent and *de novo* recurrent tumors. Comparison was performed by Wilcoxon test. (G) Differentially expressed genes of malignant cells between true recurrent and *de novo* recurrent tumors. The most significant genes with p values=0 were all shown in the plot. (H) The expression level of *GDF15* was confirmed to be higher in truly recurrent tumors than in *de novo* recurrent tumors in validation cohort 1. (I) Dotplot showing the significance and strength of specific interactions between malignant cells and DC subtypes (DC1, DC2 and mDC) in truly recurrent (green) and *de novo* recurrent samples (red). (J) Representative multiplexed immunofluorescent images showing that GDF15^+^CD45^-^ malignant cells resided near CD44^+^CD11c^+^ DCs. (K) Kaplan-Meier secondary recurrence-free survival curve in recurrent HCC patients with high or low GDF15 expression in validation cohort 1 (*n*=26 patients with 3 patients having RNA-seq data in two times of recurrence). (L) Dotplot showing the significance and strength of specific interactions between tumor-specific CD8^+^ T cells and DC subtypes (DC1, DC2 and mDC) in truly recurrent (green) and *de novo* recurrent samples (red).

Similarly, we examined the cell-cell interactions within the TIME of truly recurrent HCCs. Since the frequency of Treg, macrophages, and other immunosuppressive cells was not increased in truly recurrent HCC, we hypothesized that the differences in TIME might be associated with differences in the malignant cells. Thus, we first performed differential pathway analysis to identify the functional differences of malignant cells between two recurrent HCC types in the discovery cohort. We found that genes upregulated in HCC cells in truly recurrent tumors were enriched in immune-associated pathways such as “TNFα_signaling_via_NFkb, IFN_gamma_response and IFN_α_response”. In contrast, genes upregulated in HCC cells in *de novo* recurrent tumors belonged to the metabolism-associated pathways, including “oxidative_phosphorylation, cholesterol_homeostasis” (**Figure 6E**). Then, we further compared the functional signatures in the malignant cells (Sun et al., 2021). We found that truly recurrent tumors harbored a higher expression of “immune escape” signature, whereas *de novo* recurrent tumors exhibited a higher expression of “oxidative_phosphorylation” signature; these signatures were confirmed in samples from validation cohort 1 (**Figure 6F, Figure S5B**). These data indicate that malignant cells in truly recurrent HCC may directly mediate immune escape.

To gain additional insights into the immune escape in truly recurrent HCC, we evaluated the expression levels of immune checkpoint molecules previously reported to mediate immune-evasion (Nishino et al., 2017). The expression levels of immune checkpoint genes such as *PDCD1LG2*, *CD274*, *CTLA4,* and *VTCN1* were very low in the recurrent HCC cells (**Figure S5C**). Thus, we next analyzed the differentiated expressed genes between malignant cells of two recurrent types, and found that *GDF15* was significantly upregulated in truly recurrent malignant cells (*p*=0.000) (**Figure 6G**), which was also verified in the validation cohort 1 (**Figure 6H**). Then, we investigated the potential interactions between HCC cells and DCs, which was known to influence the antigen presentation to effector T cells (Wculek et al., 2020; Borst et al., 2020). We found a crosstalk between malignant cells and DCs via GDF15/TGFBR2, GDF15/CD48 or GDF15/CD44 only in truly recurrent HCCs (**Figure 6I**). In addition, the expression levels of *TGFBR2*, *CD48,* and *CD44* (Artz et al., 2016; Wang et al., 2021; Gao et al., 2021) were upregulated in truly recurrent HCC samples (**Figure S5D**), and our multiplexed IF data demonstrated that GDF15^+^ malignant cells resided in the proximity of CD11c^+^CD44^+^ DCs (**Figure 6J**). Functional studies have previously demonstrated that GDF15^+^ tumor cells could dampen the antitumor immunity function of DCs via GDF15/CD44 (Gao et al., 2021; Zhou et al., 2013). When we evaluated the prognostic biomarker value of GDF15 expression in malignant cells, we found that high GDF15 expression was associated with significantly shorter re-recurrence-free survival in recurrent HCC patients from validation cohort 1 (**Figure 6K**). The interactions between tumor specific CD8^+^ T cells and DCs were less frequent in truly recurrent HCC than in *de novo* ones (**Figure 6L**). Finally, we examined whether cell-cell interactions occurring in truly recurrent HCC were associated with the memory phenotype with weak cytotoxicity of CD161^+^CD8^+^ T cells. We found a crosstalk between T cell/ILCs and B cells with CD161^+^CD8^+^ T cells via the CLEC2D/KLRB1 axis only in truly recurrent HCC (**Figure S5E**). The expression of CLEC2D was detected primarily in T/NK cells and B cells (**Figure S5F, S5G**). Taken together, our data indicate that the interactions of GDF15^+^ malignant cells with DCs inhibits tumor-associated antigen presentation, hinders the activation of T cells and is associated with a poor prognosis in truly recurrent HCC.

## Discussion

Targeting TIME has emerged as an approach to preventing and treating recurrences (Zhou et al., 2017). A recent study revealed the features immune ecosystem of early relapsed, truly recurrent HCCs (Sun et al., 2021). However, differences in the immune landscape between truly and *de novo* recurrent HCC remain unclear. Resolving this issue is essential because recurrent HCCs are seldom treated based on the type of recurrence, and their TIMEs might be distinct. Indeed, our single-cell transcriptomic analyses of HBV-related recurrent HCCs demonstrated that truly recurrent HCCs were characterized by an increased frequency of CD161^+^CD8^+^ T cells with weak cytotoxicity. In contrast, *de novo* recurrent HCCs featured increased cytotoxic and exhausted CD8^+^ T cell fractions. A deeper transcriptomic profiling analysis of immunosuppressive pathways showed that truly recurrent malignant cells closely interact with DCs and T cells while myeloid cells crosstalk with T cells may mediate the their exhaustion in *de novo* recurrent HCC.

Our data demonstrate the distinct patterns of immunosuppression in the TIMEs of *de novo* versus truly recurrent HCCs (**Figure 7**). On the one hand, *de novo* malignant cells harbored higher oxidative phosphorylation capacity and infiltration by CD8^+^ T cells with cytotoxic and exhausted phenotypes. On the other hand, truly recurrent malignant cells exhibited higher immune escape capacity and upregulated GDF15 expression to dampen the antigen presentation of DCs via GDF15/CD44, which might shape the compromised anti-tumor immunity of truly recurrent HCC through enrichment in CD161^+^CD8^+^ T cells with weak cytotoxic and more quiescent phenotype. The distinct immune ecosystems of *de novo* and truly recurrent HCC emphasize the need for different immunotherapy strategies for different recurrent HCC types. For example, ICIs might be a more rational choice for *de novo* recurrent HCC, a hypothesis supported by the early clinical data in our ongoing prospective phase 2 clinical trial (TALENT; NCT04615143), which is testing the efficacy of neoadjuvant anti-PD-1 immunotherapy (tislelizumab) in resectable recurrent HCC patients after initial curative treatment (**Figure S6A**). In this trial, two achieved objective responses among the eleven patients enrolled, with one achieving a complete response and the other achieving a partial response (**Figure S6B, 6C**). Notably, the recurrent tumors of these two responders were classified as *de novo* recurrences, while those from non-responders were true recurrences. While these data are preliminary, the responses to anti-PD-1 immunotherapy in patients with *de novo* recurrent HCCs could be partially explained by our data showing cytotoxic and exhausted phenotypes of T cells and relatively normal function of DCs in their TIME. In contrast, in truly recurrent HCCs, anti-PD1 therapy might not be effective in eliciting a robust tumor-specific cytolytic attack by T cells because they reside in a resting state without efficient stimulation by tumor antigen, which may be due to the impaired antigen-presentation function by DCs and GDF15 expression in these malignant cells.

**Figure 7.**
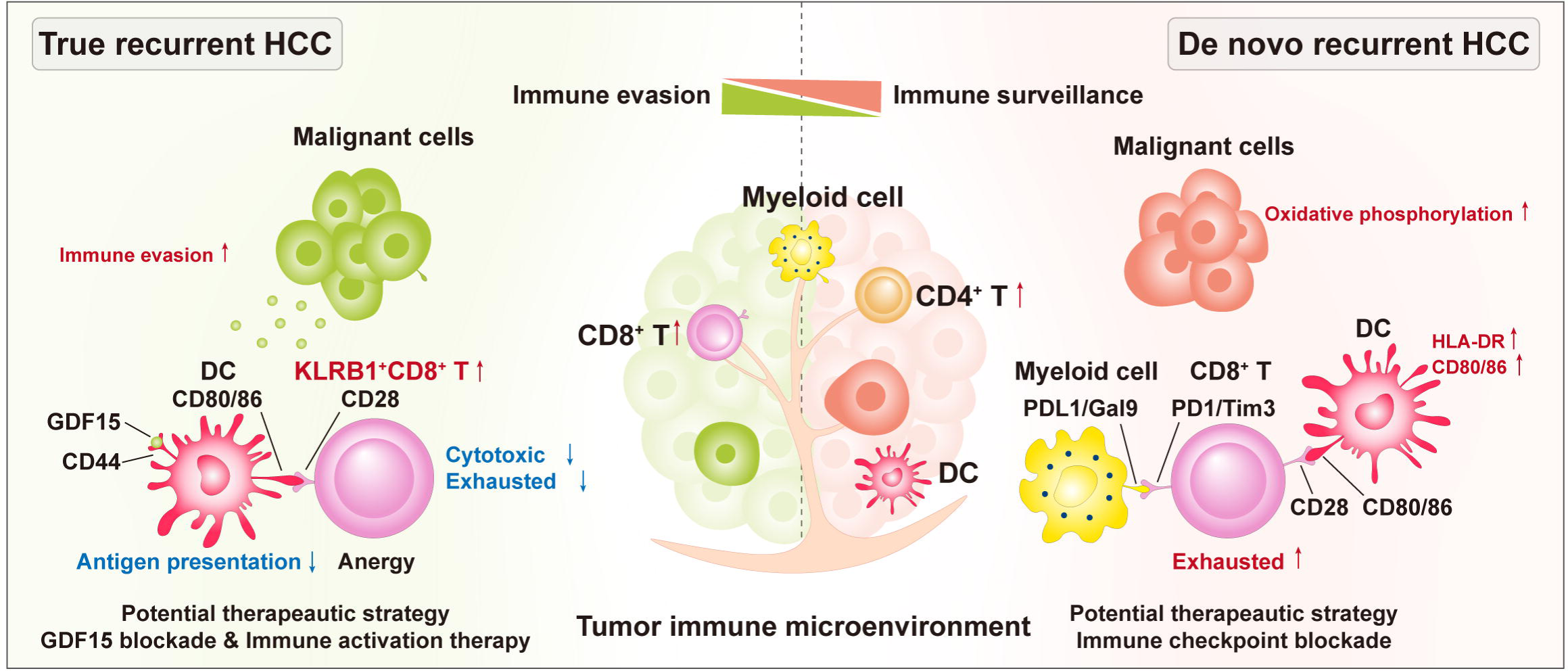
Diagram Summarizing the Distinct ScRNA-seq Immune Ecosystems between *De Novo* and Truly Recurrent HCC.

GDF15 is a member of the transforming growth factor-β (TGFβ) superfamily (Wischhusen et al., 2020). GDF15 is a potent suppressor of DCs maturation and antigen-presentation that inhibits the expression of co-stimulatory and MHC-II molecules (Gao et al., 2021; Zhou et al., 2013; Wischhusen et al., 2020). Our findings show that DC function was poorer and expression of CD80, CD86, and HLA-DR molecules was lower in truly recurrent HCC in which malignant cells overexpressed GDF15. Moreover, GDF15 can dampen immune cell trafficking and enhance the malignant cells’ stemness (Roth et al., 2010; Kempf et al., 2011; Tanno et al., 2014). Due to these immunosuppressive functions, GDF15 is a potetial biomarker of resistance to anti-CTLA-4 therapy and a target for immunotherapy (Nyakas et al., 2019; Havel et al., 2019). We discovered that GDF15^+^ malignant cells interfered with DCs only in truly recurrent HCC but not in *de novo* recurrent HCC. In addition, GDF-15 has metabolic effects associated with cancer-related cachexia (Wang et al., 2021), but whether the effect of GDF15 on cancer metabolism has any impact on DC and T cell activation remains unknown and requires future investigations. Finally, GDF15 expression correlated with patients’ survival, indicating that it might be a prognostic biomarker in truly recurrent HCCs. These data suggest that treatments targeting GDF15^+^ malignant cells or inducing neoantigen release, or helping neoantigen presentation might be more rational for truly recurrent HCC patients, for example, GDF15 blockade with or without the combination of DC vaccine or adoptive T cell transfer (Melief et al., 2015; Mardiana et al., 2019). The GDF15 neutralizing antibody CTL-002 is clinically available and is currently being tested in a phase I, first-in-human clinical trial in patients with advanced-stage solid malignancy (NCT04725474) (Melero et al., 2021).

Notably, we discovered that CD8^+^ T cells in recurrent HCC reside in different immune states, with remarkably differential transcriptional features depending on recurrence type. Specifically, CD8^+^ T cells in *de novo* recurrences displayed cytotoxic and exhausted phenotypes, whereas CD8^+^ T cells in true recurrences presented a memory phenotype with increased expression of *KLRB1*. Supporting these findings, in the transition trajectory path, CD8^+^ T cells were mainly at the pre-branch phase of cytotoxic state and the terminal end of the exhausted state in *de novo* recurrences. In contrast, the CD8^+^ T cells had considerable distribution at the fate 2 phase of relatively resting memory state in true recurrences. For truly recurrent HCC, CD8^+^ T cells had a differentiation trajectory to transit to CD8_Tem_GZMK_CD161 cells in addition to the traditional trajectory to T cell exhaustion. This differentiation trajectory might reflect a memory response in true recurrences. However, these T cells showed an upregulation in CD161 expression, a marker associated with poorer anti-tumor effector function in CD8^+^ T cells (Sun et al., 2021). CD161 exerts immunosuppressive effects by dampening the infiltration of effector T cells and cytotoxicity against malignant cells (Mathewson et al., 2021; Di et al., 2022). Our data are consistent with an immunosuppressive phenotype of CD161^+^CD8^+^ T cells in truly recurrent HCC. In contradistinction, CD8^+^ T cells could go through the traditional developmental program from cytotoxic to exhausted states in the *de novo* recurrent HCC ecosystem, consistent with a differential immune escape mechanism.

Given the distinct CD8^+^ T cell phenotypes in two types of HCC recurrence, we evaluated the transcriptomic profiles of other cells and the cell-cell interactions in the TIME. Interestingly, we found that the *de novo* recurrent malignant cells have a gene expression associated with oxidative phosphorylation and lower expression of immune escape-related genes. In the TIME of *de novo* recurrent HCCs, myeloid cells closely interacted with CD8^+^ T cells via LAGSL9/HAVCR2 or PD-L1/PDCD1 axes, which are well recognized and druggable mechanisms leading to the exhaustion of CD8^+^ T cells. On the other hand, the truly recurrent malignant cells presented more robust immune evasion capability. Our data indicate that the interaction between malignant cells and DCs and the defective antigen presentation may mediate immune evasion in truly recurrent HCCs. DCs are crucial immune cells presenting tumor antigen and play a vital role in priming anti-tumor T cell immunity (Wculek et al., 2020). In our study, we first observed that the antigen presentation capability and co-stimulatory signal of DCs were lower in truly recurrent HCCs when compared to *de novo* recurrent HCCs. Moreover, we identified that malignant cells’ crosstalk with DC1/DC2/mDC in truly recurrent HCCs via GDF15/CD44, which compromised their antigen-presentation to CD8^+^ T cells, reduced the activation of CD8^+^ T cells, and was associated with a resting memory phenotype of CD8^+^ T cells.

Interestingly, our genomic analysis of HCC recurrence type revealed that a substantial proportion of cases recurred from monoclonal seeding, implying a surprisingly early metastatic seeding (Hu et al., 2019). Moreover, a detailed analysis of the dynamics of tumor metastasis and the timing of dissemination revealed that metastasis may have happened when the primary HCC was small and not clinically detectable. In addition, genomic stratification of recurrences was inconsistent with the one used in the clinic, with a substantial fraction of *de novo* recurrent HCCs occurring within 2 years of surgical resection. Our data show that both types of recurrence may occur much earlier than currently prevailing wisdom and are relevant for the future management of recurrent HCC patients. These findings also emphasize the necessity of perioperative therapy to prevent HCC recurrence. However, none of the perioperative interventions tested so far has shown efficacy in a phase III study in unselected HCC patients, including when using effective drugs for advanced HCC (e.g., sorafenib) (Bruix et al., 2015). Our findings suggest the need to rethink perioperative therapy to prevent both true and *de novo* HCC recurrences. They also provide the rationale and a path forward to tailoring immunotherapy by considering the type of recurrence (based on genomic data) and potential targets for therapy in TIME.

In conclusion, our results reveal the different TIMEs of *de novo* and truly recurrent HBV-related HCC, as reflected in differential immune and tumor cell phenotypes, T cell developmental trajectories, TCR clonality, transcriptomic profiles, and cell-cell interactions. These data warrant similar comparative studies in HCCs of other HCC etiologies. Our findings may have immediate clinical relevance for ongoing studies of immunotherapy-based approaches in the perioperative setting in HCC, which currently lacks effective strategies. These new insights into the differential mechanisms of immunosuppression associated with the two types of recurrence also provide potential targets for more effective immunotherapy strategies to prevent and treat recurrent HCC, for example using anti-PD-1 treatment in *de novo* recurrences and GDF15 blockade or DC vaccine in true recurrences. These approaches will need to be prospectively validated in HBV-related HCC studies, ideally guided by genomic– and TIME-based characteristics for patient stratification.

## Acknowledgement

The work of the authors is supported by the National Science Fund for Distinguished Young Scholars (No. 81825013), the National Natural Science Foundation of China (No. 82172047, 81771958), the Guangdong Natural Science Foundation of Distinguished Youth Scholar (No. 2022B1515020060), the Guangdong Natural Science Foundation (No. 2021A1515010450), the Kelin Outstanding Young Scientist of the First Affiliated Hospital, Sun Yat-sen University (R08030), US National Cancer Institute grants (No. R01CA260872, R01CA260857) and DoD PRCRP Impact Award W81XWH-19-1-0284.

## Author Contribution

Conceptualization, S.L.C., D.G.D. and M.K.; Methodology, S.L.C., G.R.L., Y.B.X., H.J.H., and M.H.H.; Investigation, S.L.C, J.P.W., Z.H.D., X.X.R., X.Z.Z., Q.W.Z., G.P.Z. and C.Y.L.; Formal analysis, S.L.C., G.R.L., Y.B.X., H.J.H., M.H.H. and D.M.K.; Resource, C.H., H.C.S., W.X.X., S.L.S., S.Q.L., S.P., Q.Z. and M.K.; Data curation, M.K.; Writing-original draft, S.L.C.; Writing-review & editing, S.L.C., D.G.D. and M.K.; Visualization, S.L.C., G.R.L. and Y.B.X.; Supervision, Q.Z., D.G.D. and M.K.; Funding acquisition, S.L.C., D.G.D. and M.K.

## STAR METHODS

### SUBJECT DETAILS

#### Patient Samples and Study Design

To dissect the TIME of the two types of recurrent HCCs, we collected 23 tumor samples and 11 paired non-tumor adjacent liver tissues (NAT) from 20 treatment-naïve recurrent HCC patients (discovery cohort) and performed scRNA-seq (**Figure 1A**). In this cohort, tissue specimens, including primary tumors, NAT and matched recurrent tumors were used for WES to confirm the type of recurrence and for bulk RNA-seq to validate the findings from scRNA-seq analysis. Patient characteristics of this cohort are shown in **Table S1**. Moreover, to validate the findings in the discovery cohort, we included 26 patients with paired primary and recurrent surgical samples for WES and bulk RNA-seq (validation cohort 1), and 47 patients with paired primary and recurrent surgical samples for WES and IF (validation cohort 2) (**Figure 1B**). The baseline characteristics between patients with *de novo* recurrence and truly recurrence was compared respectively in all the three cohorts and provided in **Table S4**. The clinical follow-up was censored on Jan 31, 2022. This study was approved by the Institution Review Board of the First Affiliated Hospital of Sun Yat-sen University (Protocol No. [2020]355), Zhongshan Hospital of Fudan University, China (Protocol No. B2020-301), and Massachusetts General Hospital Boston, USA (Protocol No. 2020P000639). Signed informed consent was obtained from all patients included in the study.

Two expert pathologists confirmed the diagnosis and determined tumor content after pathological reviews. The tissues were snap-frozen in liquid nitrogen immediately within 15 minutes after resection for bulk sequencing, and fresh tissues were prepared into single cell suspension for sc-RNAseq and flow cytometry analysis.

### METHODS DETAILS

#### Exome Capture, Library Preparation and Sequencing

Total DNA was extracted from the snap-liver frozen tissues using QIAGEN DNeasy Blood & Tissue Kit (Qiagen, Hilden, Germany). The qualified genomic DNA of tissue samples was fragmented to 200-300 bp by Covaris technology with resultant library, and then adapters were ligated to both ends of the fragments.

Next, extracted DNA was amplified by ligation-mediated PCR (LM-PCR), then purified and hybridized to the Agilent human exome array for enrichment. Non-hybridized fragments were washed out. Both non-captured and captured LM-PCR products were subjected to real-time PCR to estimate the magnitude of enrichment. Each captured library was then loaded on a HiSeq X TEN platform (Illumina, San Diego, California, USA), and sequences of each individual patient were generated as 150bp paired-end reads.

For formalin-fixed paraffin embedded samples, the extracted DNA was first analyzed using agarose-gel electrophoresis to assess the data integrity and any degradation. Qualified DNA samples were randomly fragmented into 200–300 bp using Covaris technology. The exome regions were captured using Agilent human exome array. The captured DNA were then sequenced on HiSeq X TEN platform, generating 2×150bp paired-end reads.

#### Reads Mapping and Variation Detection

The adapter sequence in the raw data was removed, then reads with high quality were gapped aligned to the NCBI human reference genome (hg19) using BWA by default parameters (Li et al., 2009). We performed local realignment of the original BAM alignment using the GATK (McKenna et al., 2010) and followed by Picard to mark duplicates reads.

Somatic substitutions were detected by MuTect (Cibulskis, et al., 2013) based on BWA alignment and high confident somatic SNVs were called if the following criteria were met (I) both the tumor and normal samples should be covered sufficiently (≥ 10×) at the genomic position; (II) the variants should be supported by at least 5% of the total reads in the tumor while less than 1% in the normal; (III) the variants should be supported by at least five reads in the tumor.

High confident somatic insertions and deletions (indels) were called using the following steps: (I) candidate somatic indels were predicted with GATK Somatic Indel Detector with default parameters; (II) high confident somatic indels were defined after filtering germline events. All high confident somatic mutations were filtered out by the dbSNP (version 135) site which is commonly polymorphic without known medical impact. The remaining mutations were annotated with ANNOVAR (Wang et al., 2010) and subjected to subsequent analyses.

To increase the sensitivity of mutation calling in this study, we have applied additional procedures to identify mutations that are missed by MuTect due to low purity for the tumor sample or low coverage at specific genomic loci. Briefly, we obtain read counts for all somatic mutations across all tumor samples in the same patient. Somatic mutation was considered absent if either its variant allele frequency (VAF) was less than 0.02 or the mutation-supporting reads were fewer than 3.

#### Phylogenetic Tree Construction

We derived a mutation sequence space for each tumor, which covers all the somatic mutation sites, alternative nucleotide for mutation-positive and corresponding reference nucleotide for mutation-negative. After replacing reference nucleotides with corresponding alternative nucleotides, we obtained the mutation sequences for each tumor. Germline sample was set as an outgroup root. Sequences with 20 bp in length surrounding the somatic mutation were extracted to construct the phylogenetic trees of each patient based on the Maximum Parsimony method implemented in MEGA7 (Kumar et al., 2016). Inferred trees were then redrawn using Adobe Illustrator with relative trunk and branch lengths scaled in proportion to the number of exonic mutations.

#### Nei’s Genetic Distance

Nei’s genetic distance is widely used to assess the potential heterogeneity for stratification or substructure among individuals sampled for exploring a genetic association (Libiger et al., 2009). The recurrent tumor samples classified with the same spatial or longitudinal tag (short-term local, long-term local, long-term distant) were retained for statistical comparisons. We used the following formula to estimate Nei’s genetic distance based on cancer cell fraction (CCF) for each patient’s sample. Let x be all CCFs of sample 1 and y be all CCFs of sample 2.

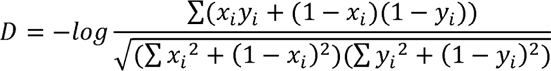

#### Tumor Purity and CCF Estimation

Tumor purity was calculated from FACETS (Shen et al., 2016) with the information of somatic mutations and copy number variations. The CCF was estimated from somatic single nucleotide variant (sSNV) frequencies and copy number profiles using PyClone (Roth et al., 2014). For each sample, the corresponding tumor purity was used to adjust the estimated CCF. Since multiple regions in the tumor tissue were sequenced in our study, a merged CCF of each sSNV was calculated to integrate the multiple region data (Equation 1).

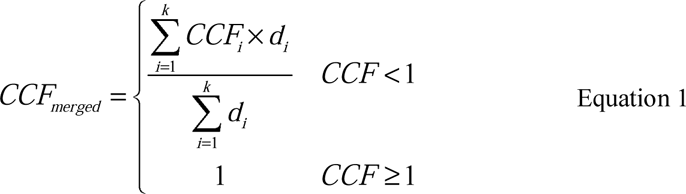

where *d_i_* and *CCF_i_* are the sequence depth and the estimated CCF value in region *i*.

#### Estimation of the Timing of Metastatic Dissemination in HCC Patients

The timing of metastatic dissemination was estimated based on the CCF of each SNV in the tumor sample. For patients with multiple tumors, the merged CCF was first calculated using the above method. According to the merged CCFs, we defined the primary private SNVs as those with a merged CCF smaller than 1% in the metastasis tumor. While, the metastasis private SNVs were similarly defined as those with merged CCF smaller than 1% in primary tumor. Then, the total number of primary-private SNVs that are present at merged CCF>10%, 20%, 40% and 60% were counted and notated as *S_1_*, *S_2_*, *S_3_* and *S_4_*, respectively. Correspondingly, the notation of *S_5_*, *S_6_*, *S_7_* and *S_8_* were also counted as the total number of metastasis-private SNVs that are present at merged CCF>10%, 20%, 40% and 60%, respectively. Besides, the number of SNVs that have merged CCF >60% in metastasis samples, and have 10%< merged CCF <60% in primary samples were counted as *S_9_*. Finally, these nine statistics were then integrated into a representation vector and inputted into SCIMET to analysis the evolutionary dynamics of a specific P/R pair.

To quantitatively estimate the timing of metastasis, SCIMET was used to model the dynamics of tumor dissemination. The patient-specific mutation rate (*u*) together with the primary tumor size (*N_d_*) at the time of dissemination were inferred from the statistical analyses described above.

#### RNA Sequencing

After total RNA was collected, poly(A) mRNA was isolated and cut into short fragments. Taking these short fragments as templates, random hexamer-primer was used to synthesize the first-strand cDNA. The second-strand cDNA was synthesized using buffer, dNTPs, RNase H and DNA polymerase I, respectively.

Short fragments were purified with QIAQuick PCR extraction kit (Qiagen, Hilden, Germany) and resolved with EB buffer for end reparation and adding poly (A). After that, the short fragments were connected with sequencing adaptors. For PCR amplification, we selected suitable fragments as templates, based on agarose gel electrophoresis. Finally, the library was sequenced using HiSeq X TEN platform, and 150bp paired-end reads were generated.

#### RNA Sequencing Data Analysis

Qualified reads were obtained after removing raw reads with adapters or of low quality and then aligned to the human genome (hg19) by HISAT (Kim et al., 2015) with default parameters. RSeQC (Wang et al., 2012) was used to measure gene expression abundance as reads per kilobase per million mapped reads (RPKM). GDF15 expression level was compared between truly and *de novo* recurrent HCC in validation cohort 1. Besides, the tumor-specific CD8^+^ T cell signature, immune escape signature and oxidative phosphorylation signature were also compared between true recurrent and *de novo* recurrent tumors in validation cohort 1. We calculated the tumor-specific CD8^+^ T cell signature score and immune escape score as the average expression of genes listed in the previous papers with ssGSEA using GSVA package (Sun et al., 2021; Simoni et al., 2018). The Wilcoxon rank sum test was performed to calculate the significance of differences between two types of recurrent HCC.

#### Multiplexed Immunofluorescence (IF)

Multiplexed IF staining was performed as previously described (Angelova et al., 2018). The 4μm-thick slides cut from the FFPE blocks were dewaxed in clarene, rehydrated by a decreasing ethanol series (100%, 95%, 70% for 5 min each) and fixed in NBF (10% neutral buffered formalin). Slides were stained to enable the simultaneous visualization of five markers– Panel 1: anti-CD8 antibody (clone ab93278, Abcam); Panel 2: anti-CD8 antibody (clone 70306S, CST), anti-CD68 antibody (clone 76437S, CST), anti-Galectin 9 antibody (clone 54330S, CST), anti-Tim3 antibody (clone 45208S, CST); Panel 3: anti-CD8 antibody (clone ab93278, Abcam), anti-CD68 antibody (clone 76437S, CST), anti-PD1 antibody (clone ab237728 Abcam), anti-PD-L1 antibody (clone 13684S, CST); Panel 4: anti-GDF15 antibody (clone ab206414, Abcam), anti-CD45 antibody (clone 13917S, CST), anti-CD44 antibody (clone ab206414, Abcam), anti-CD11c antibody (clone ab52632, Abcam)– on the same slide using PANO 7-plex IHC kit, cat 0004100100 (Panovue, Beijing, China). At each cycle of staining, microwave treatment in antigen retrieval solution pH 6.0 or pH 9.0 was applied to perform antigen retrieval. After blocking proteins for 10 min, these five primary antibodies were sequentially incubated for 60 min at room temperature, followed by the incubation of HRP-conjugated secondary antibody and tyramide signal amplification with Opal to enable their detection by fluorescence microscopy. The process was performed for the following antibodies/fluorescent dyes: Panel 1: anti-CD8/Opal-570; Panel 2: anti-CD8/Opal-690, anti-CD68/Opal-520,anti-Galectin 9/Opal-570, anti-Tim3/Opal-620; Panel 3: anti-CD8/Opal-690, anti-CD68/Opal-570, anti-PD-1/Opal-520, anti-PD-L1/Opal-620; Panel 4: anti-GDF15/Opal-620, anti-CD45/Opal-570, anti-CD44/Opal-690, anti-CD11c/Opal-520. The slides were microwave heat-treated after each TSA operation. Finally, all slides were stained with 4′-6′-diamidino-2-phenylindole (DAPI, SIGMA-ALDRICH) for 5 min.

#### Multispectral Imaging Analysis

To obtain multispectral images, the stained slides were scanned using the TissueFAXS platform (TissueGnostics, Vienna, Austria) at 10× magnification, which captures the fluorescent spectra at 20-nm wavelength intervals from 420 to 720 nm with identical exposure time; and the scans were combined to build a single stack image. Spectral libraries were established from the extracted images in which images of unstained and single-stained slides were applied to extract the spectrum of autofluorescence of tissues and fluorescein, respectively. The library was then used to unmix the multispectral images with the StrataQuest software (TissueGnostics, Vienna, Austria). Using this spectral library, reconstructed images of slides with the autofluorescence removed were acquired for imaging analysis. For each primary antibody, the cut-off value for positivity was determined according to the staining pattern and intensities of all images, respectively.

#### Single Cell Reverse Transcription, Amplification, and Sequencing

For scRNA-seq, tumor tissues were processed with mechanical chopping with blades, followed by Tumor Dissociation Kit (130-095-929, Miltenyi Biotec) and DNaseI (DN25-100MG, Sigma Aldrich) digestion in medium (RPMI1640 with 5% FBS) for 30 min at 37°C. The cell suspension was passed through a 70-um filter (130-098-462, Miltenyi Biotec), followed by 1×RBC lysis buffer (00-4333-57, eBioscience™) treatment. Single cells were processed with the GemCode Single Cell Platform using the Gemcode Gel Bead, Chip and Libraray Kits (10x Genomics, Pleasanton, CA) following the manufacturer’s protocol with modifications. Briefly, samples were processed using kits pertaining to the Chromium 5’ immune profiling v2 barcoding chemistry of 10x Genomics. Estimated 20,000 cells were loaded to each channel with the average recovery rate 8000 cells. Libraries were constructed according to the standard 10x Genomics protocol (Single Cell 5′ Reagent Kits v5.2 User Guide) with modifications and performed the paired-end 2×150bp (PE150) sequencing on Novaseq™ 6000 (llumina, San Diego, CA).

#### Single-cell RNA-seq Data Processing

The single-cell raw data were generated by 10x Genomics. Raw data were then processed by CellRanger (version 6.0.2, 10x Genomics) pipeline to obtain the feature-barcode unique molecular identifier (UMI) matrices. The human genome GRCh38 was used as reference in our study.

Before analyzing the single-cell expression data, we first removed ambient cell free mRNA contamination using SoupX (Young et al., 2020), and then performed quality control on the UMI matrices to exclude dying cells, cell fragments and doublets. The number of unique genes detected in each cell, the total number of molecules detected within a cell and the percentage of reads that map to the mitochondrial genome were calculated. We then filtered cells with unique gene counts over 6,000 or less than 200, which corresponds to doublets or cell fragments. To filter out dying cells, cells with >15% mitochondrial counts were also removed. Cells simultaneously expressed marker genes of two or more major cell types were removed in subsequent analysis.

#### Unsupervised Clustering and Cell Type Identification

We calculated the normalized expression level for each gene by diving the read counts for each cell by the total counts for that cell and multiplied by a scale factor of 1,000,000. The natural-log transformed value were taken as the final measurement of expression level for each gene in a specific cell.

Based on the normalized expression level, we next selected a subset of genes that with high cell-to-cell variation in the dataset. The top 4,000 highly variable genes were kept for downstream analysis. Then, the principal component analysis (PCA) was performed on these variable genes. Following the results of PCA, an adequate number (30-40) was determined by Elbowplot of principal components (PCs).

FastMNN correction algorithm was performed with highly variable genes as input to integrate data from different samples and correct the batch effect (Haghverdi et al., 2018). The UMap algorithm with a resolution parameter in a range of 0.1-0.7 was applied for dimensionality reduction and visualization. To identify marker genes that define a cluster, differential expression analysis was performed by comparing each single cluster to all other cells. To accelerate the computational time of differential expression analysis, genes with >0.5 log-fold difference on average between the two groups of cells and detectable expression in more than 25% of cells in either of the two groups of cells were retained. Using the above differentially expressed genes, cells were annotated to different cell types according to their well-known canonical markers. All the above analysis was performed using the Seurat R package (v 4.0) (Hao et al., 2021).

#### TCR Analysis

Single-cell V(D)J data from the 5′ Chromium 10x kit were initially processed with Cell Ranger vdj pipeline to obtain TCR contigs with alpha/beta chains. For each sample, the output file filtered_contig_annotations.csv, containing TCR α- and β-chain CDR3 nucleotide sequences, was used for downstream analysis. Only those assembled chains that were productive, highly confident, full length, with a valid cell barcode and matched alpha/beta chains were retained. If a cell had two or more qualified chains of the same type, only the chain with the highest UMI count was qualified and retained. For each, cells with an identical CDR3 amino acid sequence were considered as having originated from the same clonotype and were therefore identified as clonal cells. S_ace estimate was adopted to assess TCR richness using R package vegan (v2.5-7) (O’Hara et al., 2005).

#### Identification of Tumor-specific and Tumor-irrelevant T cells

The tumor reactive status of CD8^+^ exhausted T cells (Liu et al., 2022) and CXCL13^+^ PD-1^+^ CD4^+^ T-helper Tumor cells (Cohen et al., 2022) were first confirmed by expressing of the exclusive tumor associated exhaustion marker CXCL13 and marker genes of tumor-specific T cells including *ENTPD1*, *ITGAE*, *MAF* and *ZBED2*. Then, according to these two literatures (Liu et al., 2022, Cohen et al., 2022), T cells in the same clones with CD8_Tex_CXCL13 and CD4_Tht_CXCL13 cells, highly expressing the tumor-specific T cell signature genes including *ENTPD1*, *ITGAE*, *PDCD1*, *CXCL13*, *MAF* and *ZBED2,* were considered as tumor-specific T cells and other T cells were considered as tumor-irrelevant, and were defined together with T cells with no retained TCR contig as Unknown/irrelevant (U/I) T cells.

#### Differentially Expressed Genes Analysis in scRNA-seq Data

To define genes that may function in different types of recurrent tumors, differential expression analysis between true recurrent tumors and *de novo* recurrent tumors in specific cell groups was carried out using ‘‘FindMarkers’’ function implemented in the Seurat package. The Wilcoxon rank sum test with log-scaled fold change |logFC| > 0.6 and *P* value < 0.05 was performed to select differentially expressed genes.

#### Pathway Analysis

To reveal the potential biological functions in two types of recurrent tumors, GSEA was performed with R package ‘clusterProfiler’ between true recurrent and *de novo* recurrent tumors to identify enriched pathways in the HALLMARK gene sets released by MSigDB (Wu et al., 2021; Liberzon et al., 2011; Liberzon et al., 2015). Pathways that have a *P* value smaller than 0.05 were defined as being significantly enriched.

#### Definition and Calculation of Gene Signature Scores

To assess the functional status of CD8^+^ T cells, two functional signatures were collected from published literatures. The T cell exhausted signature contains 5 genes related to the exhaustion status of CD8^+^ T cells, and the T cell cytotoxic signature includes 12 functional genes contributed to T cell activation. Using AddModuleScore in Seurat package, the T cell exhausted score and T cell cytotoxic score were calculated for each CD8^+^ T cell. Boxplot was adopted to present the scoring difference between true recurrent tumors and *de novo* recurrent tumors, and Wilcoxon rank-sum test was performed to indicate the statistical significance.

Similarly, to measure the impact of malignant cells on immune system, the immune escape signature and oxidative phosphorylation signature were collected from previously published literatures and the HALLMARK gene sets (Sun et al., 2021; Liberzon et al., 2015). To define the functional phenotype of dendritic cell, an antigen processing and presentation signature, co-stimulation signature and DC differentiation signature were used (Liu et al., 2021). The DC co-stimulation signature was defined by 4 co-stimulatory markers, *CD40*, *CD80*, *CD86* and *CD83*.

#### Construction of Cell Developmental Trajectory

We inferred the developmental trajectory of CD8^+^ T cell using Monocle2 package (Qiu et al., 2021). The 10x Genomics sequencing data was first imported into Monocle2 using the CellDataSet class, and the negative binomial distribution was chosen to model the reads count data. Before inferring the CD8^+^ T cell trajectory, differentially expressed genes across different cell populations were selected as input features. Then, a Reversed Graph Embedding algorithm was performed to reduce the data’s dimensionality. With the expression data projected into a lower dimensional space, cells were ordered in pseudotime and trajectory was built to describe how cells transit from one state into another. After the cell trajectories were constructed, differentially expressed genes along the trajectory separated by the branch point were detected using the ‘‘differentialGeneTest’’ function. For each interested gene, the expression trend along the pseudotime was estimated using non-linear regression, and plotted with a curve chart.

#### Inference of Cell-cell Communications

R package Cellchat (v.1.1.3) was adopted to identify significant ligand-receptor pairs within true recurrent and *de novo* recurrent tumor samples. Ligand-receptor communication probabilities/strengths were computed, tested, compared and visualized on the tumor samples of truly recurrent and *de novo* recurrent HCC patients. The minimum communication cell threshold was set to 10 and other parameters were left as default.

#### Flow Cytometry Experiment and Analysis

Cryopreserved single-cell suspensions pooling from recurrent HCC tissues were thawed, washed, and blocked by Fc receptor blocking agent (564220, BD Pharmingen, San Diego, CA, USA) in staining buffer (564765, BD) for 15 min at room temperature (RT). Then suspension was labeled with Live/Dead Fixable Viability Stain (565694) to exclude dead cells. Followed by manufacturer’s instruction, cells were subsequently stained with CD45 BUV395 (563792, BD), CD14 PerCP-Cy5.5 (562692, BD), CD11c PerCP-eFluor 710 (46-0116-42, Thermo Fisher Scientific, USA), CD80 BUV563 (741404, BD), CD86 BV480 (566131, BD), CD83 BV785 (305338, Biolegend), HLA-DR BUV805 (748338, BD), CCR7 BV650 (353234, Biolegend), XCR1 BV421 (372610, Biolegend), CD1c PE-Cy7 (331516, Biolegend) in staining buffer containing Brilliant Stain Buffer (563794, BD) for 20min RT. DCs acquisition was conducted on Cytek Aurora 5 Laser Systems (Cytek Biosciences), and the collected data were further analyzed with FlowJo (Tree Star Inc.).

#### Treatment Response Evaluation for Recurrent HCC Patients Enrolled in the Phase II Clinical Trial of Anti-PD-1 Therapy

To explore treatment responses in patients with different recurrent HCC types, we evaluated the preliminary results of a prospective phase 2 clinical trial (TALENT, registered No. NCT04615143) assessing the efficacy of neoadjuvant anti-PD-1 antibody (tislelizumab) for resectable recurrent HCC patients after initial curative treatment. RECIST 1.1 was used to evaluate the treatment response of target tumors, in which complete response (CR), partial response (PR), stable disease (SD) or progressive disease (PD) were classified based on the radiologic evaluation of the target lesion. Patients with CR and PR were defined as responders, and those with SD and PD were defined as non-responders. Waterfall plot was drawn according to the immune subtypes and tumor size changes from baseline.

### STATISTICAL ANALYSIS

All data analyses were conducted in R 3.6.2. and Stata/MP 14.0. Statistical significance was defined as a two-sided *P* value of less than 0.05. We performed the comparison of gene signature scores and expression levels of marker genes between truly recurrent and *de novo* recurrent tumours using Wilcoxon rank sum test. The Nei’s genetic distance was compared by Wilcoxon rank sum test. The density of the immune cell was compared with Student’s t test. For survival analysis, we analyzed the association of GDF15 expression level and second relapse in the validation cohort 1. The samples were grouped into high and low GDF15 expression groups based on the median value. Kaplan–Meier survival curves were plotted to show differences in survival time, and log-rank p values reported by the Cox regression models implemented in the R package survival were used to determine the statistical significance.

## Supplementary Figure Legends

Figure S1. Genomic analysis identified two types of recurrence (*de novo* cancer and true recurrence) based on non-silent mutations and phylogenetic trees in the discovery cohort. (Related to data shown in Figure 1) The heat maps of non-silent mutations (left), phylogenetic trees (right) and timeline representation (bottom) of 20 patients with paired primary and recurrent HCC. Presence (orange or purple) or absence (white) of a non-silent mutation was indicated for each tumor. Driver somatic mutations were mapped to the phylogenetic trees. Recurrent tumors of the upper panel cases shared no common mutations or trunks with primary tumors, indicating their independent origin (*de novo* cancer). Recurrent tumors of the lower panel cases shared common mutations and trunks with primary tumors, indicating the true recurrence from original tumors. In *de novo* cancer type, the four cases of PT18, PT04, PT06 and PT17 recurred within 2 years.

Figure S2. **Nei’s genetic distances for groups of short-term and local recurrence, short-term and distant recurrence, and long-term and distant recurrence in discovery cohort and validation cohort 1.** Comparisons were performed between two of them using Wilcoxon rank sum test. The Nei’s genetic distance of the group of short-term and distant recurrence was not significantly different to that of the group of long-term and distant recurrence (both *p*>0.05).

Figure S3. **ScRNA-seq Landscape of Recurrent HCC Microenvironment. (Related to data shown in** Figure 3**) (A)** Bar plot showing the proportions of 23 recurrent tumor samples and 11 non-tumoral adjacent liver tissue (NAT) samples in each major cell types. (**B**) Heatmap displaying the copy number variation analysis, inferred from scRNA-seq data for tumor cells. Patient samples are labeled by different colors. (**C**) Boxplot showing the cell fraction of each major cell types in tumor and NAT samples of true recurrence and *de novo* recurrence. Comparisons were performed with Wilcoxon test. (**D**) UMAP plot showing the expression of selected marker genes for the defined myeloid cell subtypes. (**E**) Boxplot showing the cell fraction of myeloid cell subtypes in tumor and NAT samples of true recurrence and *de novo* recurrence. Comparisons were performed with Wilcoxon test. (**F**) Flow cytometry analysis of cell fraction of DC1, DC2 and mDC in truly recurrent and *de novo* recurrent tumors. Comparisons were performed with Student’s t test.

Figure S4. **Identification and Characteristics of Tumor-specific CD8^+^ T cell Subtypes in Recurrent HCC. (Related to data shown in** Figure 4**) (A)** Boxplot showing the cell fraction of CD4^+^ T cells, CD8^+^ T cells, Tregs, innate lymphoid cells and other T cells in tumor and NAT samples of true recurrence and *de novo* recurrence. Comparisons were performed with Wilcoxon test. (**B**) Boxplot showing the cell fraction of all the T cell subtypes in tumor and NAT samples of true recurrence and *de novo* recurrence. Comparisons were performed with Wilcoxon test. (**C**) Boxplot showing the cell fraction of tumor-specific CD4^+^ T cells, tumor-irrelevant CD4^+^ T cells, tumor-specific CD8^+^ T cells and tumor-irrelevant CD8^+^ T cells in tumor and NAT samples of true recurrence and *de novo* recurrence. Comparisons were performed with Wilcoxon test. (**D**) Violin plot showing the expression score of tumor-specific CD8^+^ T cell signature in truly recurrent and *de novo* recurrent tumor samples. Comparison was performed with Wilcoxon test. (**E**) UMAP plot showing the expression of selected marker genes for the defined tumor-specific CD8^+^ T cell subtypes. (**F**) GSEA enrichment of the KLRB1^+^ CD8^+^ T cells in this study for signatures of previously reported KLRB1^+^ CD8^+^ T cells. False discovery rates were determined by one-tailed permutation test by GSEA. NES, normalized enrichment score. (**G**) Alluvial plot showing the corresponding relation between functional annotations of tumor-specific CD8^+^ T cells and their general annotations in CD8^+^ T cells. (**H**) Pseudotime trajectories of CD8^+^T cell subtypes identified in panel **G** in recurrent HCC samples. (**I**) Chord plot showing the common clones shared among the five tumor-specific CD8^+^ T cell subtypes.

Figure S5. Distinct Cell-cell Interaction Patterns in *De Novo* versus Truly Recurrent HCC Microenvironments. (Related to data shown in Figure 6) (**A**) Representative multiplexed immunofluorescent images showing that PD-L1^+^CD68^+^ myeloid cells resided near PD1^+^CD8^+^ T cells. (**B**) Violin plot showing the expression score of immune escape and proliferation signatures in true recurrent and *de novo* recurrent tumor samples in bulk RNA-seq data from validation cohort 1. (**C**) Expression level of checkpoint genes in truly recurrent malignant cells and *de novo* recurrent malignant cells. Comparisons were performed with Wilcoxon test. (**D**) Violin plots showing the expression level of *TGFBR2*, *CD44* and *CD48* in DC subtypes in truly recurrent (green) and *de novo* (red) recurrent tumors. (**E**) Dotplot showing the significance and strength of CLEC2D/KLRB1, CLEC2C/KLRB1, CLEC2B/KLRB1 interactions between different cell types and tumor-specific CD8^+^ T cells in truly recurrent HCC. (**F**) UMAP plot showing the expression of *CLEC2D* in different cell subtypes. (**G**) Violin plot showing the expression level of *CLEC2D* in different cell subtypes in truly recurrent HCC.

Figure S6. **Preliminary Data from a Prospective Phase II Clinical Trial of Neoadjuvant Anti-PD-1 Immunotherapy for Resectable Recurrent HCC Patients.** (**A**) Study design of the TALENT trial of neoadjuvant anti-PD-1 antibody (tislelizumab) for resectable recurrent HCC patients after initial curative treatment. (**B**) Waterfall plot analysis for eleven enrolled patients showing maximal percentage change in the sum of the longest diameters of target lesions as compared to baseline using magnetic resonance imaging. (**C**) Pre-treatment and post-treatment magnetic resonance imaging of responder patients 1 and 2. White arrows point to the tumor lesions.

## Supplementary Table Titles

Table S1. Clinical information of the paired primary and recurrent hepatocellular carcinoma patients in discovery cohort.

Table S2. Summary of whole exome sequencing (WES) in paired primary and recurrent HCC in discovery cohort.

Table S3. The type of recurrence for patients in the validation cohort 1 and 2.

Table S4. Comparison of baseline characteristics between *de novo* recurrence and truly recurrence in three cohorts.

Table S5. The metastatic type of truly recurrent HCC patients with monoclonal seeding pattern.

Table S6. Summary of quality control data for single-cell RNA sequencing in the discovery cohort.

Table S7. Complete list of top100 differential genes in each cluster of recurrent HCC.

## References

1. Angelova, M., Mlecnik, B., Vasaturo, A., Bindea, G., Fredriksen, T., Lafontaine, L., Buttard, B., Morgand, E., Bruni, D., Jouret-Mourin, A., et al. (2018) Evolution of metastases in space and time under immune selection. Cell 175, 751–765.

2. Artz, A., Butz, S., Vestweber, D. (2016) GDF-15 inhibits integrin activation and mouse neutrophil recruitment through the ALK-5/TGF-βRII heterodimer. Blood 128, 529–541.

3. Borst, L., van der Burg, S.H., van Hall, T. (2020) The NKG2A-HLA-E axis as a novel checkpoint in the tumor microenvironment. Clin. Cancer. Res. 26, 5549–5556.

4. Bruix, J., Takayama, T., Mazzaferro, V., Chau, G.Y., Yang, J.M., Kudo, M., Cai, J.Q., Poon, R.T., Han, K.H., Tak, W.Y., et al. (2015) Adjuvant sorafenib for hepatocellular carcinoma after resection or ablation (STORM): a phase 3, randomised, double-blind, placebo-controlled trial. Lancet. Oncol. 16, 1344–1354.

5. Chen, G., Cai, Z.X., Li, Z.L., Dong, X.Q., Xu, H.P., Lin, J.L., Chen, L.H., Zhang, H.Q., Liu, X.L., Liu, J.F. (2018) Clonal evolution in long-term follow-up patients with hepatocellular carcinoma. Int J Cancer. 143, 2862–2870.

6. Chen, Y.J., Yeh, S.H., Chen, J.T., Wu, C.C., Hsu, M.T., Tsai, S.F., Chen, P.J., Lin, C.H. (2000) Chromosomal changes and clonality relationship between primary and recurrent hepatocellular carcinoma. Gastroenterology 119, 431–440.

7. Cibulskis, K., Lawrence, M.S., Carter, S.L., Sivachenko, A., Jaffe, D., Sougnez, C., Gabriel, S., Meyerson, M., Lander, E.S., Getz, G. (2013) Sensitive detection of somatic point mutations in impure and heterogeneous cancer samples. Nat. Biotechnol. 31, 213–219.

8. Cohen, M., Giladi, A., Barboy, O., Hamon, P., Li, B.G., Zada, M., Gurevich-Shapiro, A., Beccaria, C.G., David, E., Maier, B.B., et al. (2022) The interaction of CD4+ helper T cells with dendritic cells shapes the tumor microenvironment and immune checkpoint blockade response. Nat. Cancer. 3, 303–317.

9. Dahmani, A., Janelle, V., Carli, C., Richaud, M., Lamarche, C., Khalili, M., Goupil, M., Bezverbnaya, K., Bramson, J.L., Delisle, J.S. (2019) TGFβ programs central memory differentiation in ex vivo-stimulated human T cells. Cancer. Immunol. Res. 7, 1426–1439.

10. Dang, H.X., White, B.S., Foltz, S.M., Miller, C.A., Luo, J., Fields, R.C., Maher, C.A. (2017) ClonEvol: clonal ordering and visualization in cancer sequencing. Ann Oncol. 28, 3076–3082.

11. Ding, X., He, M., Chan, A.W.H., Song, Q.X., Sze, S.C., Chen, H., Man, M.K.H., Man, K., Chan, S.L., Lai, P.B.S., et al. (2019) Genomic and epigenetic features of primary and recurrent hepatocellular carcinomas. Gastroenterology. 157, 1630–1645.

12. Di, W., Fan, W.H., Wu, F., Shi, Z.F., Wang, Z.L., Yu, M.C., Zhai, Y., Chang, Y.H., Pan, C.Q., Li, G.Z., et al. (2022) Clinical characterization and immunosuppressive regulation of CD161(KLRB1) in glioma through 916 samples. Cancer Sci. 113, 756–769.

13. Duhen, T., Duhen, R., Montler, R., Moses, J., Moudgil, T., de Miranda, N.F., Goodall, C.P., Blair, T.C., Fox, B.A., McDermott, J.E., et al. (2018) Co-expression of CD39 and CD103 identifies tumor-reactive CD8 T cells in human solid tumors. Nat. Commun. 9, 2724.

14. El-Khoueiry, A.B., Sangro, B., Yau, T., Crocenzi, T.S., Kudo, M., Hsu, C., Kim, T.Y., Choo, S.P., Trojan, J., Welling Rd, T.H., et al. (2017) Nivolumab in patients with advanced hepatocellular carcinoma (CheckMate 040): an open-label, non-comparative, phase 1/2 dose escalation and expansion trial. Lancet 389, 2492–2502.

15. European Association for the Study of the Liver. (2018) EASL Clinical Practice Guidelines: Management of hepatocellular carcinoma. J Hepatol. 69, 182–236.

16. Finn, R.S., Ryoo, B.Y., Merle, P., Kudo, M., Bouattour, M., Lim, H.Y., Breder, V., Edeline, J., Chao, Y., Ogasawara, S., et al. (2020) Pembrolizumab as second-line therapy in patients with advanced hepatocellular carcinoma in KEYNOTE-240: a randomized, double-blind, phase III trial. J Clin Oncol. 38, 193–202.

17. Gao, Y.G., Xu, Y., Zhao, S.H., Qian, L.M., Song, T.T., Zheng, J., Zhang, J.F., Chen, B.L. (2021) Growth differentiation factor-15 promotes immune escape of ovarian cancer via targeting CD44 in dendritic cells. Exp. Cell. Res. 402, 112522.

18. Havel, J.J., Chowell, D., Chan, T.A. (2019) The evolving landscape of biomarkers for checkpoint inhibitor immunotherapy. Nat. Rev. Cancer. 19, 133–150.

19. Haghverdi, L., Lun, A.T.L., Morgan, M.D., Marioni, J.C. (2018) Batch effects in single-cell RNA-sequencing data are corrected by matching mutual nearest neighbors. Nat Biotechnol. 36, 421–427.

20. Hao, Y. H., Hao, S., Andersen-Nissen, E., Mauck 3rd, W.M, Zheng, S.W., Butler, A., Lee, M.J., Wilk, A.J., Darby, C., Zager, M., et al. (2021) Integrated analysis of multimodal single-cell data. Cell 184, 3573–3587.

21. Hasegawa, K., Kokudo, N., Makuuchi, M., Izumi, N., Ichida, T., Kudo, M., Ku, Y., Sakamoto, M., Nakashima, O., Matsui, O., et al. (2013) Comparison of resection and ablation for hepatocellular carcinoma: a cohort study based on a Japanese nationwide survey. J Hepatol. 58, 724–729.

22. Hu, Z., Ding, J., Ma, Z.C., Sun, R.P., Seoane, J.A., Shaffer, J.S., Suarez, C.J., Berghoff, A.S., Cremolini, C., Falcone, A., et al. (2019) Quantitative evidence for early metastatic seeding in colorectal cancer. Nat. Genet. 51, 1113–1122.

23. Imamura, H., Matsuyama, Y., Tanaka, E., Ohkubo, T., Hasegawa, K., Miyagawa, S., Sugawara, Y., Minagawa, M., Takayama, T., Kawasaki, S., et al. (2003) Risk factors contributing to early and late phase intrahepatic recurrence of hepatocellular carcinoma after hepatectomy. J Hepatol. 38, 200–207.

24. Istaces, N., Splittgerber, M., Silva, V.L., Nguyen, M., Thomas, S., Le, A., Achouri, Y., Calonne, E., Defrance, M., Fuks, F., et al. (2019) EOMES interacts with RUNX3 and BRG1 to promote innate memory cell formation through epigenetic reprogramming. Nat. Commun. 10, 3306.

25. Kempf, T., Zarbock, A., Widera, C., Butz, S., Stadtmann, A., Rossaint, J., Bolomini-Vittori, M., Korf-Klingebiel, M., Napp, L.C., Hansen, B., et al. (2011) GDF-15 is an inhibitor of leukocyte integrin activation required for survival after myocardial infarction in mice. Nat. Med. 17, 581–588.

26. Kim, D., Langmead, B. & Salzberg, S. L. (2015) HISAT: a fast spliced aligner with low memory requirements. Nat. Methods. 12, 357–360.

27. Kumar, S., Stecher, G., Tamura, K. (2016) MEGA7: Molecular evolutionary genetics analysis version 7.0 for bigger datasets. Mol Biol Evol. 33, 1870–1874.

28. Liberzon. A., Subramanian, A., Pinchback, R., Thorvaldsdottir, H., Tamayo, P., Mesirov, J.P. (2011) Molecular signatures database (MSigDB) 3.0. Bioinformatics 27, 1739–1740.

29. Liberzon, A., Birger, C., Thorvaldsdottir, H., Ghandi, M., Mesirov, J.P., Tamayo, P. (2015) The molecular signatures database (MSigDB) hallmark gene set collection. Cell. Syst. 1, 417–425.

30. Libiger, O., Nievergelt, C. M. & Schork, N. J. (2009) Comparison of genetic distance measures using human SNP genotype data. Hum. Biol. 81, 389–406.

31. Liu, B.L., Hu, X.D., Feng, K.C., Gao, R.R., Xue, Z.Q., Zhang, S.J., Zhang, Y.Y., Corse, E., Hu, Y., Han, W.D., et al. (2022) Temporal single-cell tracing reveals clonal revival and expansion of precursor exhausted T cells during anti-PD-1 therapy in lung cancer. Nat. Cancer. 3, 108–121.

32. Li, H. & Durbin, R. (2009) Fast and accurate short read alignment with Burrows-Wheeler transform. Bioinformatics 25, 1754–1760.

33. Liu, Y., He, S., Wang, X.L., Peng, W., Chen, Q.Y., Chi, D.M., Chen, J.R., Han, B.W., Lin, G.W., Li, Y.Q., et al. (2021) Tumour heterogeneity and intercellular networks of nasopharyngeal carcinoma at single cell resolution. Nat. Commun. 12, 741.

34. Llovet, J.M., Ricci, S., Mazzaferro, V., Hilgard, P., Gane, E., Blanc, J.F., de Oliveira, A.C., Santoro, A., Raoul, J.L., Forner, A., et al. (2008) Sorafenib in advanced hepatocellular carcinoma. N Engl J Med. 359, 378–390.

35. Mardiana, S., Solomon, B.J., Darcy, P.K., Beavis, P.A. (2019) Supercharging adoptive T cell therapy to overcome solid tumor–induced immunosuppression. Sci. Transl. Med. 11, eaaw2293.

36. Mathewson, N.D., Ashenberg, O., Tirosh, I., Gritsch, S., Perez, E.M., Marx, S., Jerby-Arnon, L., Chanoch-Myers, R., Hara, T., Richman, A.R., et al. (2021) Inhibitory CD161 receptor identified in glioma-infiltrating T cells by single-cell analysis. Cell 184, 1281–1298.

37. McKenna, A., Hanna, M., Banks, E., Sivachenko, A., Cibulskis, K., Kernytsky, A., Garimella, K., Altshuler, D., Gabriel, S., Daly, M., et al. (2010) The Genome Analysis Toolkit: a MapReduce framework for analyzing next-generation DNA sequencing data. Genome. Res. 20, 1297–1303.

38. Melero, I., Calvo, E., Dummer, R., Garralda, E., Schuler, M.H., Goebeler, M.E., Bargou, R.C., Gromke, T., Ramelyte, E. (2021) A phase I, first-in-human clinical trial of the GDF-15 neutralizing antibody CTL-002 in subjects with advanced-stage solid tumors (ACRONYM: GDFTHER). J. Clin. Oncol. 39, TPS2658-TPS2658.

39. Melief, C.J., van Hall, T., Arens, R., Ossendorp, F., van der Burg, S.H.. (2015) Therapeutic cancer vaccines. J. Clin. Invest. 125, 3401–3412.

40. Nishino, M., Ramaiya, N.H., Hatabu, H., Hodi, F.S. (2017) Monitoring immune-checkpoint blockade: response evaluation and biomarker development. Nat. Rev. Clin. Oncol.14, 655–668.

41. Nyakas, M., Aamadal, E., Jacobsen, K.D., Guren, T.K., Amadal, S., Hagene, K.T., Brunsvig, P., Yndestad, A., Halvorsen, B., Tasken, K.A., et al. (2019) Prognostic biomarkers for immunotherapy with ipilimumab in metastatic melanoma. Clin. Exp. Immunol. 197, 74–82.

42. O’Hara R.B. (2005) Species richness estimators: how many species can dance on the head of a pin? Journal of Animal Ecology 74, 375–386.

43. Portolani, N., Coniglio, A., Ghidoni, S., Giovanelli, M., Benetti, A., Tiberio, G.A., Giulini, S.M. (2006) Early and late recurrence after liver resection for hepatocellular carcinoma: prognostic and therapeutic implications. Ann Surg. 243, 229–235.

44. Qiu, X.J., Mao, Q., Tang, Y., Wang, L., Chawla, R., Pliner, H.A., Trapnell, C. (2017) Reversed graph embedding resolves complex single-cell trajectories. Nat. Methods. 14, 979–982.

45. Roth, A., Khattra, J., Yap, D., Wan, A., Laks, E., Biele, J., Ha, G., Aparicio, S., Bouchard-Cote, A., Shah, S.P. (2014) PyClone: statistical inference of clonal population structure in cancer. Nat. Methods. 11, 396–398.

46. Roth, P., Junker, M., Tritschler, I., Mittelbronn, M., Dombrowski, Y., Breit, S.N., Tabatabai, G., Wick, W., Weller, M., Wischhusen, J. (2010) GDF-15 contributes to proliferation and immune escape of malignant gliomas. Clin. Cancer. Res. 16, 3851–3859.

47. Ruiz, A.L., Soudja, S.M., Deceneux, C., Lauvau, G., Marie, J. (2014) NK1.1+ CD8+ T cells escape TGF-b control and contribute to early microbial pathogen response. Nat. Commun. 5, 5150.

48. Shen, R. & Seshan, V. E. (2016) FACETS: allele-specific copy number and clonal heterogeneity analysis tool for high-throughput DNA sequencing. Nucleic Acids Res. 44, e131.

49. Simoni, Y., Becht, E., Fehlings, M., Loh, C.Y., Koo, S.L., Teng, K.W., Yeong, J.P., Nahar, R., Zhang, T., Kared, H., et al. (2018) Bystander CD8+ T cells are abundant and phenotypically distinct in human tumour infiltrates. Nature 557, 575–579.

50. Sun, Y.F., Wu, L., Zhong, Y., Zhou, K.Q., Hou, Y., Wang, Z.F., Zhang, Z.F., Xie, J.R., Wang, C.Q., Chen, D.D., et al. (2021) Sing-cell landscape of the ecosystem in early-relapse hepatocellular carcinoma. Cell 184, 404–421.

51. Suzuki, R., Shimodaira, H. (2006) Pvclust: an R package for assessing the uncertainty in hierarchical clustering. Bioinformatics 22, 1540–1542.

52. Tabrizian, P., Jibara, G., Shrager, B., Schwartz, M., Roayaie, S. (2015) Recurrence of hepatocellular cancer after resection: patterns, treatments, and prognosis. Ann Surg. 261, 947–955

53. Tanno, T., Lim, Y.T., Wang, Q.J., Chesi, M., Bergsagel, P.L., Matthews, G., Johnstone, R.W., Ghosh, N., Borrello, I., Huff, C.A., et al. (2014) Growth differentiating factor 15 enhances the tumor-initiating and self-renewal potential of multiple myeloma cells. Blood 123, 725–733.

54. Villanueva, A. (2019) Hepatocellular carcinoma. N Engl J Med. 380, 1450–1462.

55. van der Leun, A.M., Thommen, D.S., Schumacher, T.N. (2020) CD8+ T cell states in human cancer: insights from single-cell analysis. Nat. Rev. Cancer. 20, 218–232.

56. Wang, D.D., Day, E.A., Townsend, L.K., Djordjevic, D., Jorgensen, S.B., Steinberg, G. (2021) GDF15: emerging biology and therapeutic applications for obesity and cardiometabolic disease. Nat. Rev. Endocrinol. 17, 592–607.

57. Wang, K., Li, M. & Hakonarson, H. (2010) ANNOVAR: functional annotation of genetic variants from high-throughput sequencing data. Nucleic Acids Res. 38, e164.

58. Wang, L., Wang, S. & Li, W. (2012) RSeQC: quality control of RNA-seq experiments. Bioinformatics 28, 2184–2185.

59. Wang, Z.W., He, L., Li, W.N., Xu, C.Y., Zhang, J.Y., Wang, D.S., Dou, K.F., Zhuang, R., Jin, B.Q., Zhang, W., et al. (2021) GDF15 induces immunosuppression via CD48 on regulatory T cells in hepatocellular carcinoma. J. Immunother. Cancer. 9, e002787.

60. Wculek, S.K., Cueto, F.J., Mujal, A.M., Melero, I., Krummel, M.F., Sancho, D. (2020) Dendritic cells in cancer immunology and immunotherapy. Nat. Rev. Immunol. 20, 7–24.

61. Wei, Y., Lao, X.M., Xiao, X., Wang, X.Y., Wu, Z.J., Zeng, Q.H., Wu, C.Y., Wu, R.Q., Chen, Z.X., Zheng, L.M., et al. (2019) Plasma cell polarization to the immunoglobulin G phenotype in hepatocellular carcinoma involves epigenetic alterations and promotes hepatoma progression in mice. Gastroenterology. 156, 1890–1904.

62. Wischhusen, J., Melero, I., Fridman, W.H. (2020) Growth/differentiation factor-15 (GDF15): from biomarker to novel targetable immune checkpoint. Front. Immunol. 11, 951.

63. Wu, S.Z., AI-Eryani, G., Roden, D.L., Junankar, S., Harvey, K., Andersson, A., Thennavan, A., Wang, C.F., Torpy, J.R., Bartonicek, N., et al. (2021) A single-cell and spatially resolved atlas of human breast cancers. Nat. Genetics. 53, 1334–1347.

64. Wu, T.Z., Hu, E.Q., Xu, S.B., Chen, M.J., Guo, P.F., Dai, Z.H., Feng, T.Z., Zhou, L., Tang, W.L., Zhan, L., et al. (2021) clusterProfiler 4.0: A universal enrichment tool for interpreting omics data. Innovation (NY) 2: 100141.

65. Xiao, X., Lao, X.M., Chen, M.M., Liu, R.X., Wei, Y., Ouyang, F.Z., Chen, D.P., Zhao, X.Y., Zhao, Q.Y., Li, X.F., et al. (2016) PD-1^hi^ identifies a novel regulatory B-cell population in human hepatoma that promotes disease progression. Cancer. Discov. 6, 546–559.

66. Young, M.D., Behjati, S. SoupX removes ambient RNA contamination from droplet-based single-cell RNA sequencing data. GigaScience, Volume 9, Issue 12, December 2020.

67. Zheng, C.H., Zheng, L., Yoo, J.K., Guo, H., Zhang, Y., Guo, X., Kang, B., Hu, R., Huang, J.Y., Zhang, Q., et al. (2017) Landscape of infiltrating T cells in liver cancer revealed by single-cell sequencing. Cell 169, 1342–1356.

68. Zhang, Q.M., He, Y., Luo, N., Patel, S.J., Han, Y., Gao, R., Modak, M., Carotta, S., Haslinger, C., Kind, D., et al. (2019) Landscape and dynamics of single immune cells in hepatocellular carcinoma. Cell 179, 829–845.

69. Zhang, Y.Y., Chen, H.Y., Mo, H.N., Hu, X.D., Gao, R.R., Zhao, Y.H., Liu, B.L., Niu, L.J., Sun, X.Y., Yu, X., et al. (2021) Sing-cell analyses reveal key immune cell subsets associated with response to PD-L1 blockade in tripe-negative breast cancer. Cancer Cell. 39, 1578–1593.e8.

70. Zhou, G., Sprengers, D., Boor, P.P.C., Doukas, M., Schutz, H., Mancham, S., Pedroza-Gonzalez, A., Polak, W.G., de Jonge, J., Gaspersz, M., et al. (2017) Antibodies against immune checkpoint molecules restore functions of tumor-infiltrating T Cells in hepatocellular carcinomas. Gastroenterology. 153, 1107–1119.

71. Zhou, Z., Li, W.L., Song, Y., Wang, L.L., Zhang, K., Yang, J., Zhang, W., Su, H.C., Zhang, Y.Q. (2013) Growth differentiation factor-15 suppresses maturation and function of dendritic cells and inhibits tumor-specific immune response. PLoS. ONE. 8, e78618.

