## Supplementary figures and images for "Distinct Single-cell Immune Ecosystems Distinguish True and *De Novo* HBV-related Hepatocellular Carcinoma Recurrences"

### Figure S1

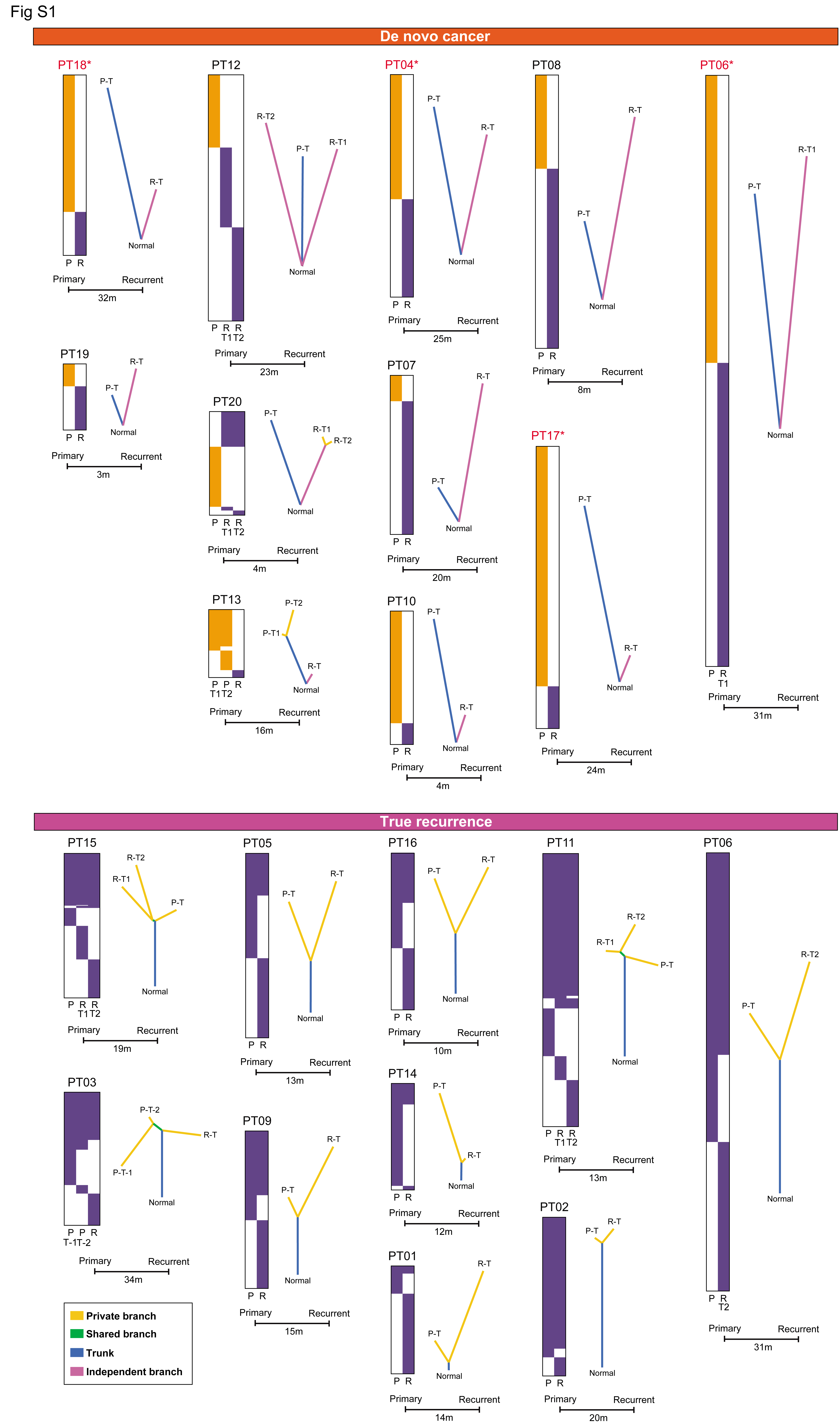

### Figure S2

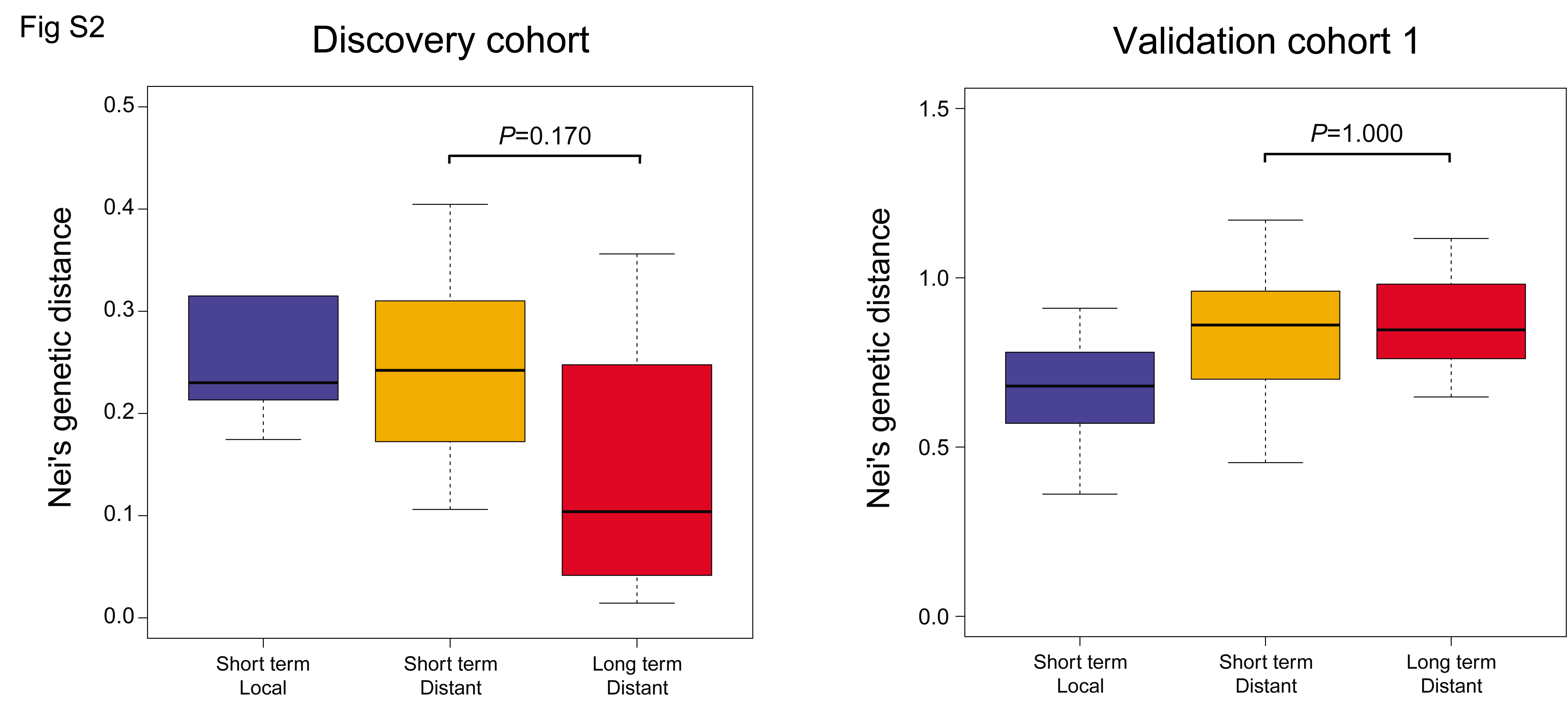

### Figure S3

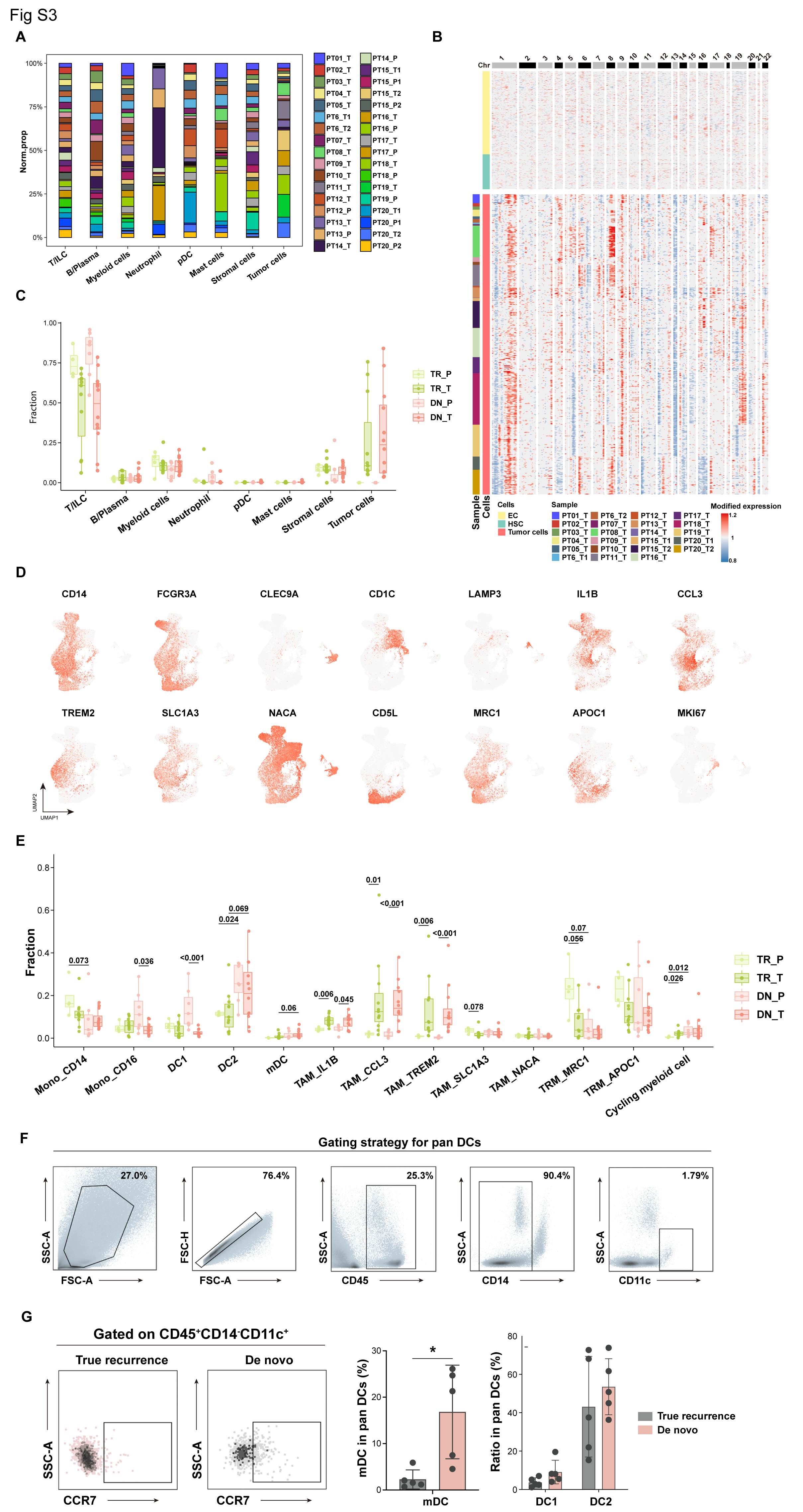

### Figure S4

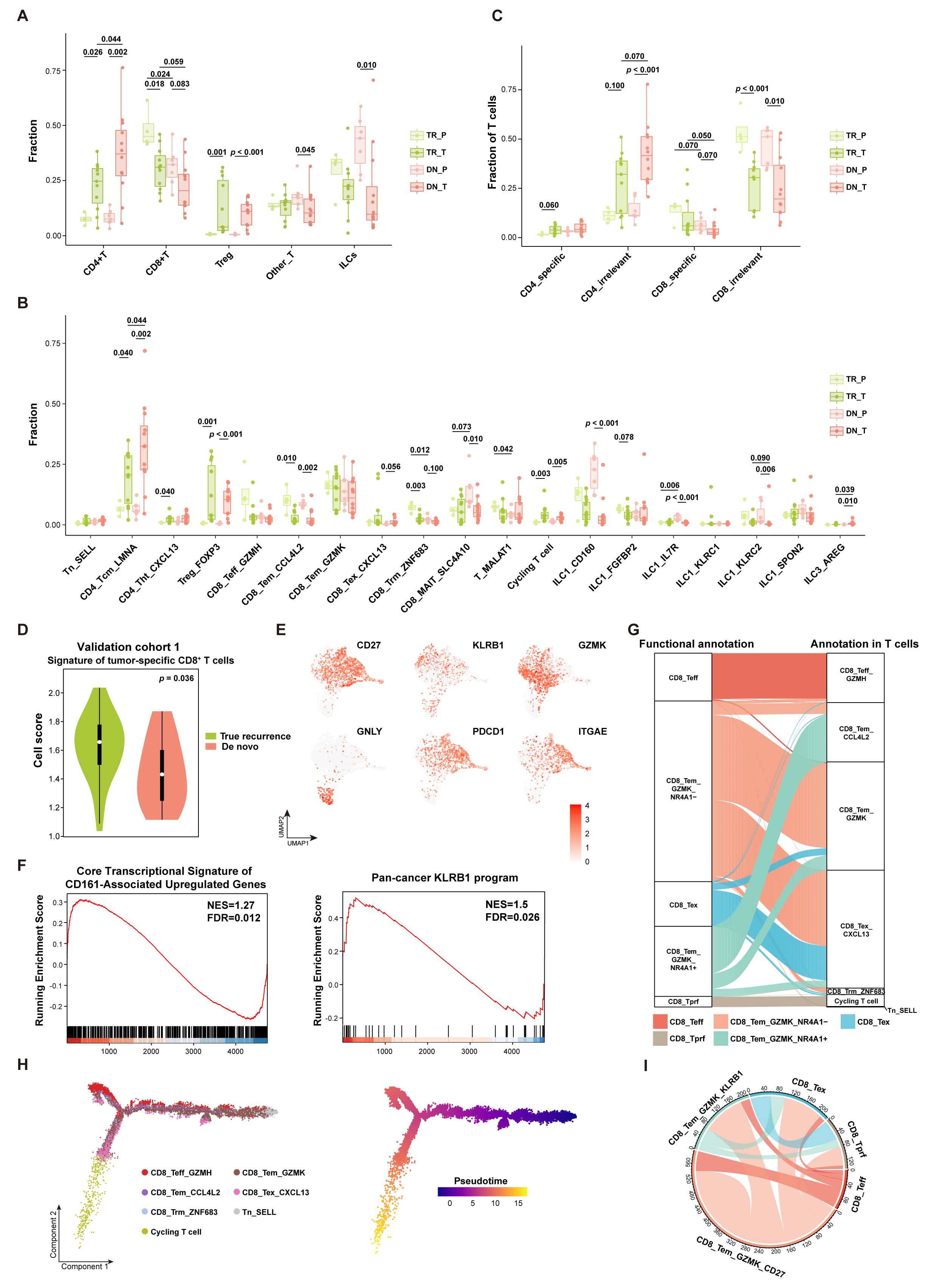

### Figure S5

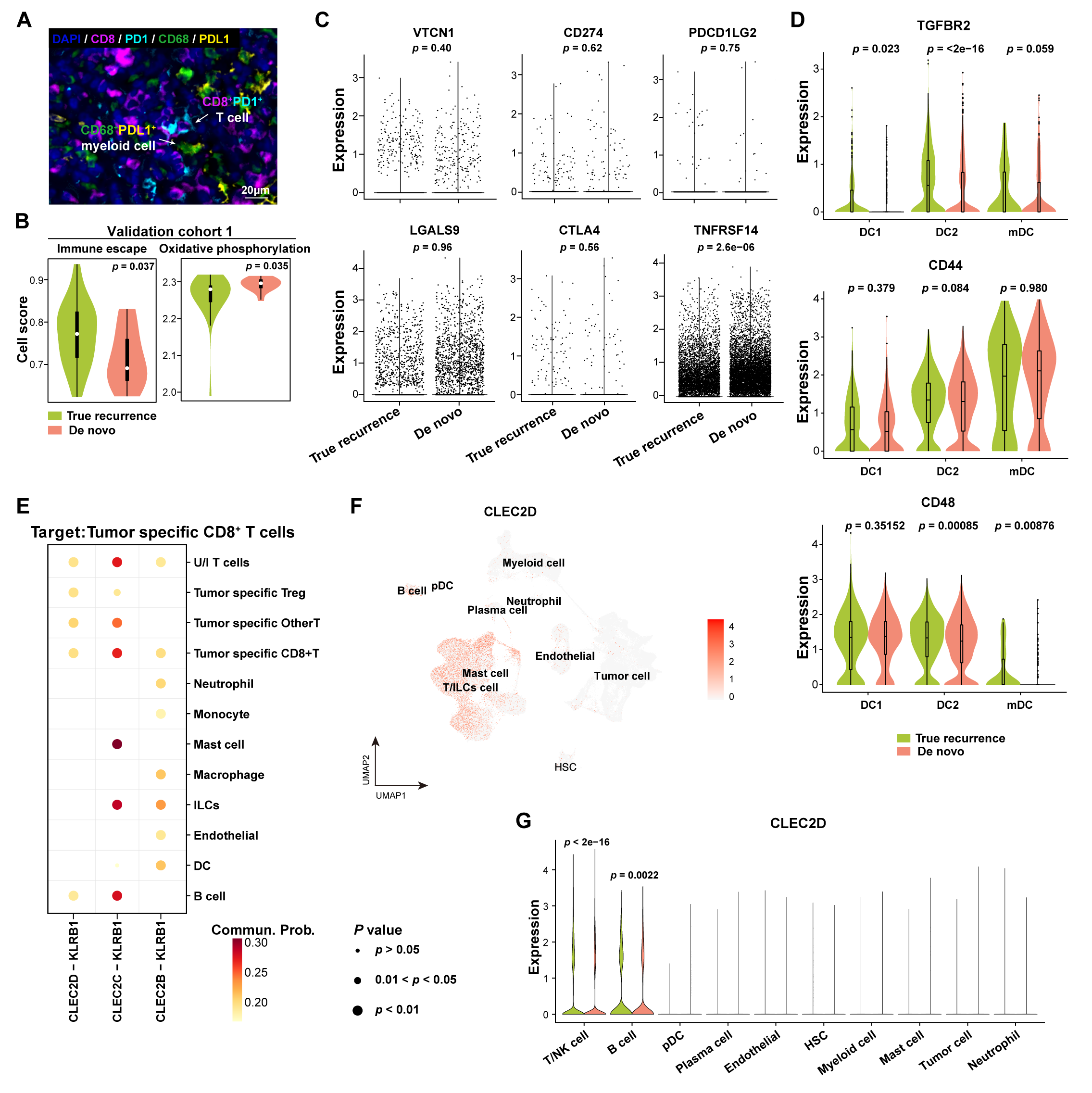

### Figure S6

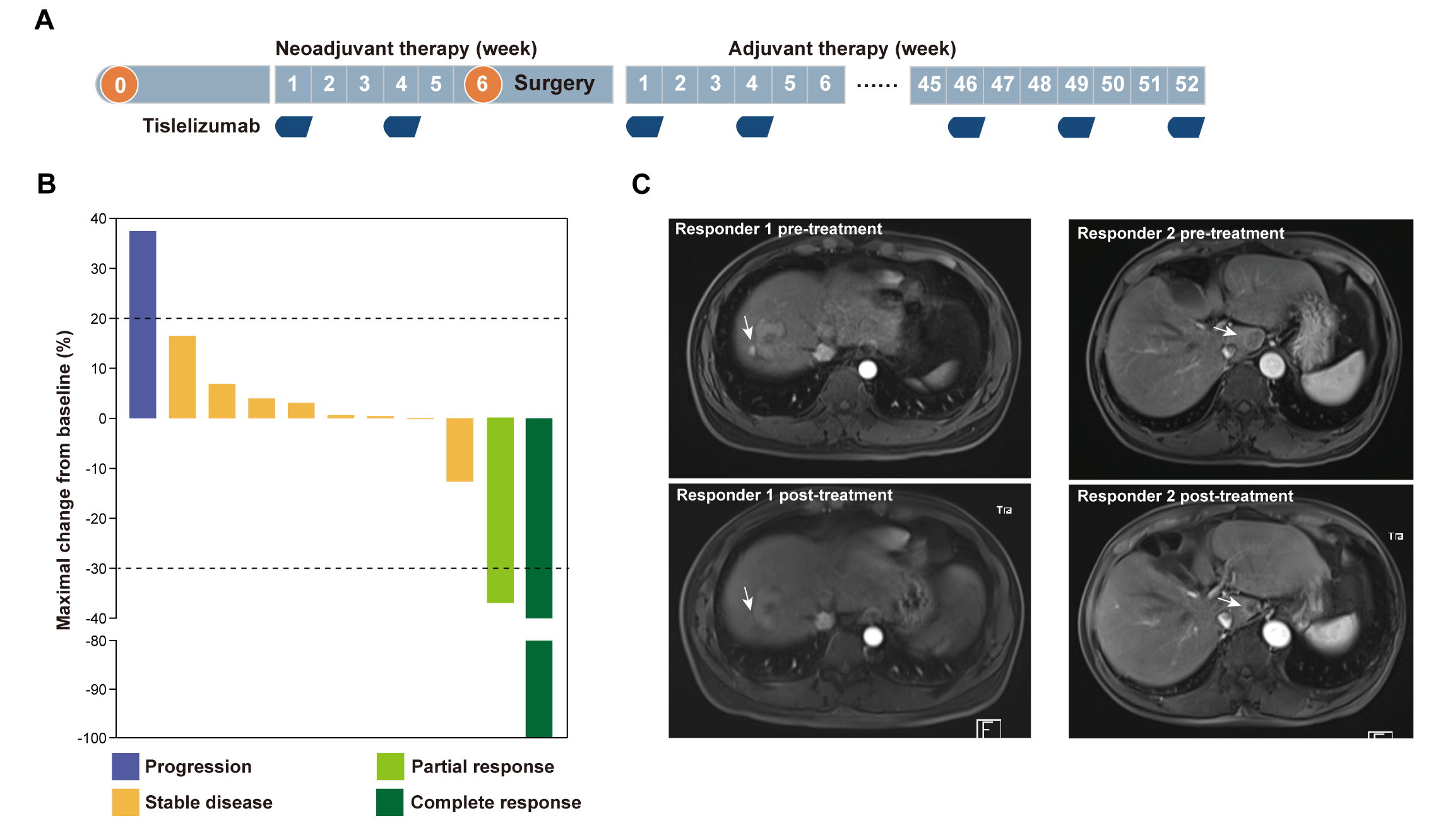
